# Development of the ULK1-Recruiting Chimeras (ULKRECs) to enable proximity-induced and ULK1-dependent degradation of mitochondria

**DOI:** 10.1101/2024.04.15.589474

**Authors:** Niyaz Zaman, Natasha Aley, Valeria Pingitore, David L Selwood, Robin Ketteler

**Author notes:** To whom correspondence should be addressed: Robin Ketteler.

## Abstract

Targeted protein degradation (TPD) has opened new opportunities to investigate signalling pathways as a research tool, and as a unique therapeutic strategy using bifunctional chimeric small molecules, with candidate molecules in clinical trials for the treatment of breast cancer and prostate cancer. Most current TPD approaches use the 26S proteasomal machinery via PROteolysis TArgeting Chimeras (PROTACs), however, new emerging strategies using the autophagy system, termed AUtophagy TArgeting Chimeras (AUTACs) expand on the degrader arsenal and repertoire of targets that can be degraded. This includes non-protein molecules such as lipid droplets, organelles, insoluble protein aggregates as well as typical TPD targets, soluble intracellular proteins. AUTACs were proposed to operate by binding the target of interest (TOI) and linking it to an autophagy cargo protein (LC3 or p62), tethering the TOI into forming autophagosomes. In this study, we designed an alternative strategy for AUTACs, reasoning that the local recruitment and activation of ULK1 is sufficient to induce the formation of an autophagosome at the site of recruitment. As a proof of concept, we used an ULK1 agonist linked to a mitochondrial targeting ligand and termed these chimeric molecules ULK1-Recruiting Chimeras (ULKRECs). We show that local activation of ULK1 by ULKRECs at the outer mitochondrial membrane (OMM) induces mitophagy, further enhanced by mitochondrial insult. Using Parkinson’s disease (PD) patient-derived fibroblasts, we show the ULKRECs induce mitophagy independently of the PRKN/PINK axis, components required to signal for canonical mitophagy in response to stressors and often dysfunctional in many neurological diseases. We propose that ULKRECs are a novel class of degraders that have potential as unique therapeutics for diseases where dysfunctional mitophagy plays a key role in disease pathology and progression.

## INTRODUCTION

Targeted manipulation of autophagy offers multiple opportunities for new discoveries into the regulation of cell biological processes and the development of therapeutic strategies. To date, very few such tools exist. Genetic interference with key autophagy genes has proven very useful to dissect the molecular basis of autophagy, identify potential drug targets and understand the role of autophagy in development and disease^1–3^. However, genetic modifications are technically challenging and time-consuming. On the other hand, chemical approaches involving small molecule inhibitors and activators operate on a much shorter time scale and modulate autophagy in an acute, reversible manner. Compounds have been developed to target a selected number of a few key proteins in the pathway such as, p62/SQSTM1 inhibitors^4^, ULK1/2 inhibitors^5,6^ and activators^7,8^ and ATG4B inhibitors^9–11^. There are several limitations in small molecule discovery, notably the inability to target what is considered the non-druggable proteome and an inability to direct the modulation locally.

An alternative is the use of targeted protein degradation (TPD). TPD has attracted a lot of attention, both as research tools for pathway modulation as well as for therapeutic strategies^12^. Most approaches to date utilise the ability of the proteasome to degrade single proteins via proximity induced lysine mono- or poly-ubiquitination of the target, marking the protein of interest (POI) for degradation by the 26S proteasome. This can be achieved using a heterobifunctional chimeric small molecule linking a targeting ligand specific to a POI with an E3 ubiquitin ligase binder, which is often referred to as a PROteasome TArgeting Chimera, or PROTAC^13^. Since its inception in the early 2000s, the method has been continuously refined, and currently, over 100 proteins have been successfully targeted using this approach (see PROTACpedia database https://protacpedia.weizmann.ac.il/ptcb/main) with more than 10 in clinical trials^14^. A recent alternative to degradation via the proteasome is the facilitated degradation of a target of interest (TOI) by the autophagosome/lysosome system. Some examples of AUtophagy TArgeting Chimeras (AUTACs) have been described, enabling the degradation of organelles such as mitochondria and non-protein molecules such as lipid droplets, as well as soluble intracellular proteins and insoluble protein aggregates^15–19^. Current strategies for AUTACs focus on proximity of the target to the autophagosome, which can be facilitated by using a small molecule binder of autophagosome membrane associated proteins such as MAP1LC3B (LC3) or LC3 interacting proteins, for example p62, in the chimeric molecule, linked to a targeting ligand^15,17,19,20^. A recent study indicated that LC3 mediated AUTACs (also referred to Autophagy Tethering Chimeras (ATTECs)) may in fact have a proteasomal mechanism, the details of the proposed mechanisms will require further study^21^. A second, significantly less mechanistically understood strategy relies on the selective pseudo-S-guanylation of a TOI to mark it for degradation in an autolysosome^16^.

Instead of bringing the target in proximity to the autophagosome or marking it for degradation, we propose that the initiation of an autophagosome at the target site is an alternative approach to achieve targeted autophagic degradation. The initiation of autophagosomes is largely controlled by a complex network of protein kinases of which ULK1/2 are central. Activation of the ULK1/2 kinases results in the activation of a downstream cascade that triggers the activation of another kinase, PI3K class III or vps34^22^, and subsequent reactions of lipid transfer via ATG2/WIPI proteins^23^. This is followed by membrane remodelling via ATG9^24^, and conjugation of the autophagosome with the ubiquitin-like proteins of the ATG8 family via a complex conjugation machinery involving ATG5, 12, 16L1, 3, 7, 10 and 4^25^. It has been shown that recruitment of ULK1/2 to specific locations in the cell can initiate the formation of an autophagosome at those sites^26^, using a chemically inducible dimerization assay involving cells stably expressing 2xFKBP translational fusions of GFP-tagged ULK1 and FRB-Fis1, a mitochondrial fission protein found on the outer mitochondrial membrane (OMM). However, these tools require the transfection of the genetic constructs, and are not amenable to acute modulation of the pathway, nor suitable for therapeutic development, but chiefly, demonstrate the principle of localised activation of autophagy. In this study we have generated the first novel heterobifunctional chimeric chemical entities that direct the ULK1 kinase to the target. Using an ULK1 agonist linked to a mitochondrial targeting ligand, we demonstrate the induction and selective degradation of mitochondria upon treatment of cells with these chimeric molecules. To recognize the ability of these molecules to target mitochondria to the autophagosome via recruitment of ULK1, we termed these AUTAC molecules, ULKRECs.

## MATERIALS AND METHODS

### Cell Culture and Chemicals

HEK293T (human embryonic kidney) (CRL-3216), PANC-1 human pancreatic ductal adenocarcinoma (PDAC) (CRL-1496) and SH-SY5Y human neuroblastoma (CRL-2266) cells were purchased from ATCC. ULK1/2 WT MEF (SV40) (RRID:CVCL_5A51)^27^, ULK1+ULK2 DKO MEF (SV40) (RRID:CVCL_5A57)^27^ were gifted by Professor Sharon Tooze (The Francis Crick Institute, London, United Kingdom). HEK293T, PANC-1 ULK1/2 WT MEF and ULK1+ULK2 DKO MEF cells were cultured in High Glucose DMEM + GlutaMAX^TM^ + Pyruvate (ThermoFisher Scientific), supplemented with 10% foetal bovine serum (FBS) and 1% penicillin/streptomycin at 37°C/5% CO_2_ (standard conditions). SH-SY5Y were cultured in High Glucose DMEM + GlutaMAX^TM^ (ThermoFisher Scientific), supplemented with 10% FBS and 1% penicillin/streptomycin under standard conditions. For mito-mKeima experiments, SH-SY5Y cells are cultured in the same media without phenol red. SH-SY5Y expressing mito-mKeima were produced as previously described^28,29^. The wild-type control human fibroblasts and PD-PINK1 null culture containing a frameshift deletion in PINK1 (c.261_276del16;p.Y90L fsx12) were acquired from the UCL BioBank under Human Tissue Act Licence Number 12198, provided by UCL Queen Square Institute of Neurology, University College London. The primary fibroblasts were cultured in High-glucose DMEM supplemented with 10% heat inactivated FBS, 1% penicillin/streptomycin and 0.2 mM uridine. Cells were split at 80% confluency and media was refreshed every 2-3 days. Torin 1 (#475991) and Bafilomycin A1 (#B1793) were purchased from Sigma-Aldrich, dissolved in dimethyl sulfoxide (DMSO) and stored at -20°C, Antimycin (#A8674) and Oligomycin (#75351) were purchased from Sigma-Aldrich and stored at -20°C. BL-918 (#S0819) and LYN-1604 (#S8597) were purchased from SelleckChem, dissolved in DMSO and stored at -80°C. AUTAC4 (#HY-134640) was purchased from MedChemExpress, dissolved in DMSO and stored at -80°C. NZ-65, NZ-66 were synthesised in house (See *Chemistry*), dissolved in DMSO and stored at -80°C or in lyophilised forms for long-term storage.

### Lentiviral production

2.5x10^5^ HEK293T cells were seeded into a 6-well plate. The subsequent day, cells were co-transfected with 900 ng psPAX2 (psPAX2 was a gift from Didier Trono (Addgene plasmid #12260; http://n2t.net/addgene:12260; RRID:Addgene_12260)), 100 ng pMD2.G (pMD2.G was a gift from Didier Trono (Addgene plasmid #12259; http://n2t.net/addgene:12259; RRID:Addgene_12259)) and 1μg of pDEST-CMV mCherry-GFP-LC3B WT (Addgene plasmid # 123230; http://n2t.net/addgene:123230; RRID:Addgene_123230, Agrotis & Ketteler)^30^ per well using X-tremeGene HP (Roche, #6366244001) at 2 μl/μg of total DNA in antibiotic-free and serum-free Opti-MEM^TM^ media (ThermoFisher). At 19 hours post-transfection, medium was replaced with complete DMEM. Viral supernatant was harvested at 48 hours post-transfection, filtered through a 0.22 μm low protein-binding syringe filter and stored at -80°C

### mCherry-EGFP-LC3 Tandem Reporter

PANC-1 cells were seeded at a medium density in 24-well plates and infected overnight with 500 μL of filtered viral supernatant containing polybrene at a final concentration of 8 μg/mL. Cells were selected at 48 hours post-transduction with 1 μg/mL puromycin and expanded into 10 cm tissue-culture treated dishes. Puromycin treatment was withheld once control cells had died. Experiments were performed without puromycin present, using pooled populations of transduced cells. PANC-1 stably expressing mCherry-EGFP-LC3 tandem reporter cells were imaged with the PerkinElmer Opera Phenix or Opera Phenix Plus High Content Imaging Systems and analysed using Columbus software (v2.9.1). Briefly, Hoechst 33342 was used to identify nuclei followed by determining regions of interest corresponding to cytoplasm for both EGFP and mCherry channels. This was followed by calculating the relative fluorescent intensity (RFI) of EGFP and mCherry for their respective cytoplasm. Spots were identified and gated for intensity greater than mean RSI for each channel under negative control conditions. Autophagosomes were defined as having EGFP and mCherry fluorescence. Autolysosomes were defined as only having mCherry fluorescence. Excitation and emission wavelengths for each channel are as follows: Hoechst 33342 = 405/435-80 nm, EGFP = 488/500-550 nm, mCherry = 561/570-630 nm, respectively. Images captured with a water immersion lens at 40x magnification.

### Western Blots

Cells were homogenised in protein lysis buffer consisting of 0.1 M Tris pH 8.0, 0.1 M NaCl, 5% glycerol, IGEPAL® CA-630 (#I8896. Sigma-Aldrich), 5 mM EDTA, 0.1 M N-ethyl maleimide (ThermoFisher, #23030) and protease inhibitor (Cell Signalling Technologies, #5871). Protein concentrations were determined using the BCA assay (ThermoFisher) to ensure equal loading. After addition of loading buffer, protein samples were electrophoresed at 100 V for 75 minutes and transferred to Bio-Rad Trans-blot Turbo PVDF membranes using the Bio-Rad turbo-blot system. PVDF membranes were subsequently blocked with TBS with 0.05% Tween-20 (#P1379, Sigma-Aldrich) (TBS-T) + 3% bovine serum albumin (BSA) (#A7906, Sigma-Aldrich). The membranes were immunoblotted with primary antibodies, overnight at 4°C. ULK1 (D8H5) Rabbit mAb (#0854) [1:1000], Phospho-ULK1 (Ser 317) Rabbit antibody (#37762) [1:1000], LC3B Rabbit antibody (#2775) [1:1000], TOM20 (D8T4N) Rabbit mAb (#42406) [1:1000] and, Phospho-Ubiquitin (Ser 65) (E2J6T) Rabbit mAb (#62082) [1:500] were obtained from Cell Signalling Technologies. Recombinant Anti-Vinculin Rabbit antibody (EPR8185) (#ab129002) [1:5000] and Anti-Mitofusin2 mouse antibody (6A8) (#ab56889) [1:1000] were purchased from Abcam. Anti-β-Actin mouse monoclonal antibody (#A1978) [1:1000] and Anti-rabbit p62/SQSTM1 (#P0067) [1:1000], were purchased from Sigma-Aldrich. Mono- and polyubiquitinylated conjugates monoclonal antibody (FK2) (BML-PW8810-0100) [1:500] was purchased from Enzo Life Sciences. Membranes were then incubated with horseradish peroxidase (HRP)-conjugated secondary antibodies (Rabbit #7074 and Mouse #7076) purchased from Cell Signalling Technologies for 1 hour and the HRP signal was detected using EZ-ECL Enhanced Chemiluminescence Detection Kit for HRP (#20-500-500, Sartorius), before imaging with an Alliance Q9 Advanced (UVITec). Where required, membranes were stripped with mild stripping buffer consisting of 0.2 M glycine, 3.47 mM SDS, 1% Tween 20, pH 2.2, blocked for 1 hour with TBS-T + 3% BSA and re-probed with primary antibodies.

### Imaging and measuring mitophagy using mito-mKeima

SH-SY5Y stably expressing mito-mKeima were produced as previously described^28,29^. On day 1, cells were seeded in 384 well CellCarrier Ultra plates (PerkinElmer) in complete media without phenol red. On day 2, cells were treated with increasing concentrations of BL-918, AUTAC4, NZ-65 or NZ-66 24 hours prior to treatment with increasing concentrations of mitochondrial uncouplers to cause mitochondrial insult on Day 3. SH-SY5Y mitochondria were treated with DMSO, 3 μM CCCP, 10 μM CCCP, 0.1/0.1 μM antimycin/oligomycin (AO), 0.5/0.5 μM AO or 1/1 μM AO. Plates were imaged over an 18-hour period at hourly intervals, immediately after the addition of mitochondrial uncouplers using a PerkinElmer Opera Phenix Plus HCS microscope and analysed using Columbus software. Hoechst 33342 dye was used to identify cell nuclei and mito-mKeima green was used to define the cytoplasm. The spot detection feature in Columbus was then used to identify the number of spots and calculated the area of the spots in the mito-mKeima green channel and mito-mKeima red channel in each cell. Spots are gated on their intensity being greater than the mean intensity for each respective channel under DMSO treated conditions. The mitophagy index was defined as the ratio of the total mito-mKeima red spot area to the total mito-mKeima green spot area. A higher mitophagy index is indicative of higher mitophagic activity. Excitation and emission spectra for each channel used were as follows: Hoechst 33342 = 405/435-480 nm, mito-mKeima green = 488/650-760 nm, mito-mKeima red = 561/570-630 nm, respectively. Images were captured with a water immersion lens at 63x magnification.

### Tetramethylrhodamine methyl ester (TMRM) assay

SH-SY5Y cells were seeded in a 96 well PerkinElmer PhenoPlate 24h prior to treatment with compounds. The following day, cells were incubated for 30 minutes with 3.24 µM Hoechst 33342 (#H3570, ThermoFisher), 100 nM TMRM and 20 nM MitoTracker Green (ThermoFisher Scientific). Cells were then treated with compounds, including 10 μM CCCP as a positive control before immediately imaging with a PerkinElmer Opera Phenix Plus High Content Screening system. Images were captured over the course of 18 hours and TMRM intensity in mitochondrial regions was quantified over time. TMRM intensity is directly proportional to changes in mitochondrial membrane potential (ΔΨ_m_). Excitation and emission spectra for each channel are as follows: Hoechst 33342 = 405nm/435-480 nm, MitoTracker Green = 488/500-550 nm and TMRM = 640/690-720 nm.

### Immunofluorescence

For TOM20/ULK1 immunofluorescence, cells were seeded in a 384 well PerkinElmer CellCarrier Ultra plate, 24 hours prior to treatment with compounds. Cells were treated for 18 hours after which they were fixed with 4% paraformaldehyde (PFA) in phosphate buffered saline (PBS) and blocked with 10% FBS and 0.1% Triton in PBS. Plates were incubated at 4°C overnight, with 1:400 mouse anti-TOM20 (Santa Cruz, #sc-17764) and 1:400 rabbit anti-ULK1 (Cell Signalling Technologies, #0854). The following day, plates were incubated with 1:1000 goat anti-mouse Alexa 488 (ThermoFisher Scientific, #A-11029), 1:1000 goat anti-rabbit Alexa 568 (ThermoFisher Scientific, #A-11011) and 1.62 µM Hoechst 33342 at room temperature for 1 hour before imaging. For ATP5a/LC3 immunofluorescence, cells were fixed with ice-cold 1:1 MeOH:acetone and blocked with 3% BSA (Sigma-Aldrich, #A3803) in Dulbecco’s PBS (D-PBS). Plates were incubated overnight at 4°C with 1:400 LC3B (D11) XP Rabbit mAb Alexa Fluor 647 Conjugate (Cell Signalling Technologies, #65299) and 1:400 ATP5a (Cell Signalling Technologies, #18023) lightning-linked to Alexa Fluor 488. The following day, plates were incubated with 1.62 µM Hoechst 33342 at room temperature for 1 hour before imaging. Excitation and emission spectra for each channel are as follows: Hoechst 33342 = 405/435-480 nm, Alexa 488 = 488/500-550 nm and Alexa 568 = 561/570-630 nm. Images were acquired at 40x and 63x magnification with water immersion lens.

### Plasmid Transfection and Imaging

pMRX-IP/Venus-mULK1 was a gift from Noboru Mizushima (Addgene plasmid #58743; http://n2t.net/addgene:58743; RRID:Addgene_58743)^31^. PANC-1 cells were seeded in antibiotic-free, serum-free Opti-MEM media (ThermoFisher Scientific) in a PerkinElmer 96-well PhenoPlate before transfecting with 100ng Venus-mULK1 in lipofectamine 3000 (ThermoFisher Scientific). 24 hours post-transfection, transfection media was replaced with normal culture media and cells were incubated for a further 24 hours. After a total transfection of 48 hours under standard conditions (see *Cell Culture*), compounds were added and subsequent image capture and/or recordings were acquired over 18 hours. 30 minutes prior to imaging, cells were incubated with 3.24 µM Hoechst 33342 and 25 nM MitoTracker Deep Red (ThermoFisher Scientific). Images were captured using the PerkinElmer Opera Phenix Plus High Content Imaging System at 63x magnification with water immersion lens. Excitation and emission spectra for each channel used are as follows: Hoechst 33342 = 405/435-80 nm, Venus = 488/515-550 nm, MitoTracker Deep Red = 640/690-720 nm.

### Computational Modelling

The X-ray crystal structure of TSPO (PDB code: 2MGY) and the kinase domain of ULK1 (PDB code: 4WNP) were downloaded from the Protein DataBank (PDB). A set of conformers were generated for each ligand using the ETKDG method, which were each energy minimised using a Merck Molecular Force Field (MMFF), using RDKit in Python 3.8. Both proteins were prepared as standard, briefly, hydrogens were added, hydrogen bonds were optimised, and protonation states of histidine residues were assigned. For ULK1, the previously described putative binding pocket consisting of Arg18, Lys50, Asn86 and Tyr89^8^ was used to define the grid for docking. For TSPO, the binding pocket occupied by PK11195 in 2MGY was used to define the grid for docking. Conformational screening was performed using GOLD Docking and scored with the ChemPLP scoring function^32^ and protein-ligand interactions were analysed using BIOVIA Discovery Studio.

### Chemistry

All commercially available solvents and reagents were used without further treatment as received unless otherwise noted. Solvent evaporations were performed under reduced pressure (40-60 °C) by using a Buchi Rotavapor R-210. Reactions under an inert and anhydrous atmosphere were performed using commercial nitrogen. Thin layer chromatography (TLC) was performed with qualitative purposes on aluminium silica gel plates (Alugram Sil G/UV 254) with detection by UV light (λ 254 nm) charring with p-anisaldehyde, KMnO4, ninhydrin, phosphomolybdic acid or with cerium ammonium molybdate reagent ((NH_4_)6Mo7O_24_·4H_2_O; Ce(SO_4_)_2_ in a 1:10 mixture of concentrated H_2_SO_4._ in H_2_O). Column purifications were carried out in a Biotage Isolera Four Flash Chromatography System using the appropriate chromatography column (normal phase Biotage Sfär Silica 5g/10g/25g and reverse phase Biotage Sfar C18 D-Duo 100 Å 12g/30g/60g). 1H-and 13C-NMR spectra were performed at the UCL Chemistry NMR Facility with Bruker DRX 500 or 600 MHz spectrometers. Chemical shifts (δ) are expressed in ppm relative to TMS and coupling constants (J) in Hz. DMSO-*d_6_*, CDCl_3_ and CD_3_OD were used as solvents at room temperature except when indicated. Chemical shifts are calibrated using residual solvent signals (DMSO-*d_6_*: δ(H) = 2.50, δ(C) = 39.52; CDCl_3_: δ(H) = 7.26, δ(C) = 77.16; CD_3_OD: δ(H) = 3.31, δ(C) = 49.00). NMR multiplicity abbreviations as follows: s = singlet, d = doublet, t = triplet, q = quartet, p = pentet, h = hextet; examples of complex multiplicities, dt = doublet of triplets, tdd = triplet of doublets of doublets. High-resolution mass spectrometry (HRMS) was performed via electron spray ionisation (ESI) using an ASAP-HESI ionisation connected to the Q Exactive Plus mass spectrometer, and were performed at UCL Chemistry Mass Spectrometry Facility. LC-MS spectra were obtained using a single quadrupole LC/MSD XT mass spectrometer with electrospray ionisation (ESI), using an analytical C18 column (Kinetex 5 µm 100 Å, 50 x 4.6 mm) and a gradient 10% → 95% MeCN + 0.1% formic acid (FA) in H_2_O + 0.1% FA (6.5 min). Solvent abbreviations: H_2_O = Water, DCM = dichloromethane, MeCN = acetonitrile, THF = tetrahydrofuran, Et_2_O = diethyl ether, EtOAc = ethyl acetate, Cy = Cyclohexane.

#### 2-oxo-2-(2-phenyl-1H-indol-3-yl)acetyl chloride

Synthesis has been previously described^32^ and was followed accordingly. Briefly, oxalyl chloride (0.620 mL, 7.22 mmol) was added dropwise to a stirring mixture of 2-phenylindole (1.2 g, 6.2 mmol) in dry Et_2_O (20 mL) at 0°C and stirred at room temperature for 4 hours. The resulting green precipitate was collected by vacuum filtration to obtain the desired product (1.6 g, 5.6 mmol, 90%) as a green solid.

#### *N*-(2-(2-(2-azidoethoxy)ethoxy)ethyl)-2-oxo-2-(2-phenyl-1H-indol-3-yl)acetamide (1)

Azido-PEG2-Amine (150 mg, 0.9 mmol) was dissolved in dry DCM (3 mL), the solution was cooled at 0°C and NaHCO_3_ (70 mg, 0.8 mmol) was added followed by 2-oxo-2-(2-phenyl-1*H*-indol-3-yl)acetyl chloride (200 mg, 0.7 mmol). After stirring at RT for 1 hour, the solvent was removed under reduced pressure and the crude product was dissolved in EtOAc, washed with H_2_O, dried over MgSO_4_, filtered, and concentrated under reduced pressure. The crude product was purified by flash chromatography column on silica gel (EtOAc 50% in Cy) to yield the title compound (150 mg, 0.4 mmol, 57%) as a green oil.

^1^H-NMR (500MHz, MeOD, *J* Hz) δ ppm 8.99 (s, 1H), 8.18 (d, 1H, *J* = 7.70), 7.42 (q, 3H, *J* = 6.44), 7.30 (quintet, 5H *J* = 7.29), 6.80 (t, 1H, *J* = 5.75), 3.25 (t, 2H, *J* = 6.90), 3.11 (q, 2H, *J* = 6.77), 1.57 (quintet, 2H, *J* = 7.15), 1.46 (t, 2H, *J* = 7.33), 1.39 – 1.21 (m, 4H).

^13^C-NMR (500MHz, MeOD) δ ppm 186.87, 163.81, 147.44, 135.69, 132.46, 129.64, 129.08, 128.71, 128.21, 124.09, 123.10, 121.98, 111.37, 110.64, 51.48, 39.37, 29.81, 29.20, 28.84, 26.53, 26.49.

HRMS (ESI-MS) m/z calc. for C_22_H_24_N_5_O_4_ [M+H] ^+^ = 422.17500, found 422.18228

#### *N*-(6-azidohexyl)-2-oxo-2-(2-phenyl-1H-indol-3-yl)acetamide (2)

6-Azido-hexylamine (0.06 mL, 4.3 mmol) was dissolved in dry DCM (1 mL), the solution was cooled at 0°C and NaHCO_3_ (35 mg, 0.4 mmol) was added followed by 2-oxo-2-(2-phenyl-1*H*-indol-3-yl)acetyl chloride (95 mg, 0.3 mmol). After stirring at RT for 1 h, the solvent was removed under reduced pressure and the crude material was dissolved in EtOAc, washed with H_2_O, dried over MgSO_4_, filtered, and concentrated under reduced pressure. The crude product was purified by flash chromatography column on silica gel (EtOAc 50% in Cyclohexane) to yield the title compound (77 mg, 0.2 mmol, 67%) as a green oil.

^1^H-NMR (500MHz, MeOD, *J* Hz) δ ppm 10.21 (s, 1H), 8.17 (d, 1H, *J* = 8.00), 7.29 – 7.1 (m, 6H), 6.92 (dd, 3H, *J* = 7.45, 21.81), 3.66 – 3.56 (m, 6H), 3.43 (t, 2H, *J* = 5.10), 3.35 (t, 2H, *J* = 4.83), 3.22 (q, 2H, *J* = 5.12).

^13^C-NMR (500MHz, MeOD) δ ppm 186.72, 171.31, 164.76, 147.76, 136.05, 132.05, 129.37, 129.11, 128.95, 128.51, 128.27, 123.78, 122.93, 121.57, 112.14, 110.17, 77.43, 77.17, 76.92, 70.71, 70.50, 70.22, 69.31, 60.53, 50.78, 39.24, 27.05, 21.14, 14.31.

HRMS (ESI-MS) m/z calc. For C_22_H_23_N_5_O_2_ [M+H] ^+^ = 390.18518, found 390.19245

#### (R)-4-(1-amino-2-((2,4-difluorophenyl)amino)-2-oxoethyl)phenyl isobutyl carbonate

To a solution of Boc-(*R*)-2-amino-2-(4-hydroxyphenyl)acetic acid (1.9 g, 7.3 mmol) in dry THF (25 mL) at -20°C, *N*-Methylmorpholine (1.6 mL, 15 mmol) and isobutyl chloroformate (1.9 mL, 15 mmol) were sequentially added. After stirring for 30 minutes at -20°C, 2,4-difluoroaniline (0.9 mL, 9.3 mmol) was added and the mixture was stirred overnight at RT. The solvent was removed under reduced pressure and the crude material was diluted with DCM, washed with 1 M HCl, 1 M NaHCO_3_, and H_2_O. The organic layers were dried over MgSO₄, filtered concentrated under reduced pressure, purified by C18 reverse phase chromatography (5% to 95% MeCN + 0.1% formic acid in H_2_O + 0.1% formic acid) and dissolved in 4 M HCl in 1,4-dioxane (5 mL). After stirring at RT overnight, the solvent was removed under reduced pressure obtaining the corresponding deprotected amine as a white solid (450 mg, 1.2 mmol, 16%, 2 steps).

^1^H-NMR (500MHz, CDCl_3_, *J* Hz) δ ppm 7.828 – 7.736 (m, 1H), 7.689 (d, 2H, *J* = 8.223), 7.353 (d, 2H, *J* = 8.637), 7.070 – 7.006 (m, 1H), 6.975 (d, 1H, *J* = 9.240), 5.325 (s, 1H), 4.035 (d, 2H, *J* = 6.547), 2.053 – 1.990 (m, 1H), 0.988 (d, 6H, *J* = 6.744).

^13^C-NMR (500MHz, CDCl_3_) δ ppm 167.70, 154.88, 153.98, 131.76, 131.02, 130.97, 127.53, 127.45, 123.39, 112.31, 112.28, 112.13, 112.10, 105.31, 105.12, 105.10, 104.91, 76.05, 57.80, 57.48, 29.02, 19.11.

HRMS (ESI-MS) m/z calc. for C_19_H_21_N_2_F_2_O_4_ [M+H] ^+^ = 379.13911, found 379.14639

#### (*R*)-2-(3-(3,5-bis(trifluoromethyl)phenyl)ureido)-N-(2,4-difluorophenyl)-2-(4-hydroxyphenyl)acetamide

The previous product (160 mg, 0.42 mmol) was dissolved in dry MeCN (5 mL) and stirred with DIPEA (1 mL, 6 mmol) for 10 minutes at RT. 3,5-Bis(trifluoromethyl)phenyl isocyanate (0.2 mL, 1.2 mmol) was added dropwise, and the reaction mixture was stirred overnight at 50°C. The solvent was removed under reduced pressure and the crude material was diluted with DCM, washed with 1 M HCl, 1 M NaHCO_3_, and H_2_O. The organic layers were dried over MgSO₄, filtered and concentrated under reduced pressure and used directly in the next step. The crude product (50 mg, 0.08 mmol) was dissolved in THF (1 mL) and cooled to 0°C before adding 2 M NaOH (1 mL) to hydrolyse the carbonate ester. After stirring at RT overnight, the solvent was removed under reduced pressure and the crude material was diluted with EtOAc, washed with 1 M HCl, 1 M NaHCO_3_, and H_2_O, dried over MgSO₄, filtered, and concentrated under reduced pressure. The crude product was purified by C18 reverse phase chromatography (5% to 95% MeCN + 0.1% formic acid in H_2_O + 0.1% formic acid) obtaining the desired product (35 mg, 0.07 mmol, 17%, 2 steps) as an off-white solid.

^1^H-NMR (600MHz, MeOD, *J* Hz) δ ppm 7.99 (s, 2H), 7.73 (td, 1H, *J* = 8.89, 5.97), 7.47 (s, 1H), 7.36, (d, 1H, *J* = 8.55), 6.985 (ddd, 1H, *J* = 10.763, 8.763, 2.828), 6.905 (dddd, 1H, *J* = 9.245, 8.061, 2.855, 1.439), 6.82 (d, 1H, J = 8.55), 5.51 (s, 1H).

^13^C-NMR (600MHz, MeOD) δ ppm 172.44, 162.20, 160.57, 158.88, 157.38, 156.29, 155.72, 143.02, 133.20, 129.79, 129.49, 127.51, 125.64, 123.84, 122.99, 118.86, 116.62, 115.62, 111.99, 104.90, 58.63.

HRMS (ESI-MS) m/z calc. for C_23_H_16_F_8_N_3_O3 [M+H] ^+^ = 534.09857, found 534.10584

#### (*R*)-2-(3-(3,5-bis(trifluoromethyl)phenyl)ureido)-*N*-(2,4-difluorophenyl)-2-(4-((1-(2-(2-(2-(2-oxo-2-(2-phenyl-1*H*-indol-3-yl)acetamido)ethoxy)ethoxy)ethyl)-1*H*-1,2,3-triazol-4-yl)methoxy)phenyl)acetamide [NZ-65] & (*R*)-2-(3-(3,5-bis(trifluoromethyl)phenyl)ureido)-i*N*-(2,4-difluorophenyl)-2-(4-((1-(6-(2-oxo-2-(2-phenyl-1*H*-indol-3-yl)acetamido)hexyl)-1*H*-1,2,3-triazol-4-yl)methoxy)phenyl)acetamide [NZ-66]

(*R*)-2-(3-(3,5-bis(trifluoromethyl)phenyl)ureido)-*N*-(2,4-difluorophenyl)-2-(4-hydroxyphenyl)acetamide (35 mg, 0.07 mmol) was dissolved in dry MeCN and anhydrous K_2_CO_3_ (36 mg, 0.26 mmol) was added. After stirring for 10 minutes at RT, 80% propargyl bromide in toluene (0.025 mL, 0.33 mmol) was added and stirred at 80°C for 48 hours. The solvent was removed under reduced pressure and the crude material was diluted with EtOAc, washed with 1 M HCl, 1 M NaHCO_3_, and H_2_O, dried over MgSO₄ filtered and concentrated under reduced pressure. The crude alkyne product (**3**) was dissolved in a mixture of THF:H_2_O = 10:1 (1 mL) followed by compound **1** or **2** (6 mg, 0.01 mmol), CuSO_4_ (0.4 mg, 0.0025 mmol) and sodium ascorbate (1 mg, 0.005 mmol). After stirring at 50°C overnight, the solvent was removed under reduced pressure and the crude product was purified by chromatography column on silica gel (5% MeOH in DCM) obtaining the final ULKRECs **NZ-65** (10 mg, 0.010 mmol, 14%, 2 steps) and **NZ-66** (11 mg, 0.011 mmol, 16%, 2 steps) as a yellow oil and yellow solid respectively.

**NZ-65** ^1^H NMR (600 MHz, DMSO-*d_6_*, *J* Hz) δ 12.38 (s, 1H), 8.53 (d, 1H, J = 5.4), 8.19 (q, 1H, J = 7.1), 8.08 (d, 1H, J = 7.4), 7.55 (dd, 2H, J = 4.2, 2.1), 7.52 – 7.41 (m, 5H), 7.32 (q, 1H, J = 9.5), 7.24 (dt, 2H, J = 21.8, 7.4), 7.06 (d, 1H, J = 10.6), 6.80 (d, 1H, J = 12.6), 6.64 (s, 1H), 5.63 – 5.54 (m, 1H), 5.11 (dd, 1H, J = 27.9, 12.7), 4.52 (p, 1H, J = 7.3), 4.37 (d, 1H, J = 8.6), 4.34 – 4.30 (m, 1H), 4.25 (q, 1H, J = 6.1), 4.18 (ddd, 2H, J = 19.6, 9.2, 6.4), 3.83 (tdd, 3H, J = 12.9, 9.5, 5.2), 3.50 (dt, 1H, J = 7.4, 3.6), 3.42 (s, 2H), 3.38 (s, 4H), 3.28 (s, 1H), 3.18 (s, 2H), 2.88 (h, 2H, J = 6.0).

^13^C NMR (600 MHz, DMSO-*d_6_*) δ ppm 187.54, 176.70, 175.76, 175.70, 172.87, 170.90, 170.04, 166.66, 147.48, 135.78, 131.35, 129.63, 129.34, 128.07, 127.35, 123.36, 122.32, 120.97, 111.99, 109.27, 105.64, 105.46, 105.29, 93.12, 91.35, 88.23, 87.95, 87.92, 76.01, 75.08, 74.86, 74.83, 73.28, 73.13, 73.04, 72.22, 69.76, 69.44, 68.70, 68.22, 66.41, 50.13, 49.97, 49.46, 48.63, 40.01, 38.16, 37.88, 37.58.

HRMS (ESI-MS) m/z calc. For C_48_H_41_F_8_N_8_O_7_ [M + H] ^+^ = 993.88423, found 993.29650.

**NZ-66** ^1^H NMR (600 MHz, DMSO-*d_6_*, *J* Hz) δ 12.37 (s, 1H), 8.44 (d, 1H, J = 5.7), 8.23 (dd, 1H, J = 19.1, 4.1), 8.09 (d, 1H, J = 7.7), 7.95 (dd, 1H, J = 12.9, 10.5), 7.81 – 7.71 (m, 1H), 7.55 (d, 2H, J = 4.1), 7.49 – 7.41 (m, 5H), 7.33 (ddd, 1H, J = 13.9, 8.0, 3.1), 7.25 (dt, 2H, J = 21.4, 7.5), 7.07 (d, 1H, J = 8.8), 5.89 (d, 1H, J = 4.7), 5.31 (q, 1H, J = 3.0), 4.35 (t, 1H, J = 7.1), 3.73 (dddt, 2H, J = 15.4, 7.8, 5.6, 3.6), 2.71 (s, 2H), 2.42 (t, 1H, J = 8.1), 1.91 – 1.62 (m, 11H), 1.17 (s, 4H).

^13^C NMR (600 MHz, DMSO-*d_6_*) δ ppm 203.53, 203.20, 187.85, 177.91, 169.67, 166.45, 157.82, 153.92, 147.40, 135.79, 131.38, 129.62, 129.31, 128.05, 127.40, 123.34, 122.29, 120.97, 111.98, 109.31, 103.06, 101.88, 98.89, 97.18, 68.31, 66.40, 66.31, 66.14, 66.05, 65.73, 65.58, 52.17, 49.36, 40.96, 40.74, 40.24, 40.10, 40.04, 38.18, 37.58, 33.17, 32.01, 31.86, 31.75, 29.66, 28.10, 27.45, 25.82, 25.55, 23.33, 23.07, 23.00, 22.87, 22.19, 21.78.

HRMS (ESI-MS) m/z calc. for C_48_H_41_F_8_N_8_O_5_ [M+H] ^+^ = 961.88623, found 961.30667

## RESULTS

To improve autophagosome-targeted degradation, we aimed to develop chimeric molecules that initiate autophagosome biogenesis at the cellular location of the cargo. We reasoned that chimeric small-molecule recruitment of ULK1 to a substrate and subsequent activation of ULK1 may be sufficient to induce the formation of an autophagosome. Previously described genetic constructs have shown that local activation of ULK1 is sufficient for local autophagosome biogenesis^26^. Therefore, we were interested in exploring compounds that bind to and activate ULK1 as the autophagy inducer. We assessed currently known ULK1 agonists as potential autophagy inducers for the design of ULKRECs. We tested the ability of two published ULK1 agonists, LYN-1604^7^ and BL-918^8^ to activate autophagy using the tandem mCherry-EGFP-LC3 reporter. Briefly, the tandem reporter exhibits EGFP and mCherry fluorescence in autophagosomes, and only mCherry fluorescence in autolysosomes as the EGFP is quenched due to the low pH of the autolysosomes^30^. We observed a significant increase in autophagosome and autolysosome number upon treatment with BL-918, supporting its role as an activator of autophagy (Figure 1A-C). To support these observations, we also demonstrated an induction of phosphorylation of ULK1 at Ser317, an activating phosphorylation, upon treatment with BL-918 (Figure 1D-E) and a reduction in p62/SQSTM1 levels (Figure 1F) with concomitant increases in the LC3I:LC3II ratio (Figure 1G), further supporting its role as an ULK1 agonist and autophagy activator. We did not see any consistent effects on autophagy by treatment with LYN-1604. We also assessed the mitophagic potential of BL-918. Under baseline conditions, we found BL-918 was unable to induce degradation of TOM20, suggesting it is unable to induce mitophagy itself under baseline conditions (Supplementary Figure 1A, B). We co-validated this using SH-SY5Y cells stably expressing the mito-mKeima reporter. Under neutral pH (for example, in the cytoplasm), mito-mKeima is excited in the green fluorescence wavelength range, while in acidic conditions (as observed in lysosomes), it is excited in the red fluorescence wavelength range. We used the ratio of total red over green puncta area to derive a mitophagy index, directly proportional to mitophagic activity. The mito-mKeima assay supports the prior observation, BL-918 does not induce mitophagy under baseline conditions (Supplementary Figure 1C). Furthermore, we also find that at experimentally relevant concentrations, BL-918 does not enhance mitophagy in response to mitochondrial toxins, CCCP or antimycin/oligomycin. We found that BL-918 does enhance mitophagy at concentrations greater than or equal to 20 µM, when co-treating with mitochondrial toxins (Supplementary Figure 1D, E), however, we attribute this to a depolarisation of the mitochondrial membrane potential at these higher concentrations (Supplementary Figure 1F), using the TMRM assay as previously described^33,34^ therefore, the observed mitophagy is an artefact of mitochondrial stress. Overall, we confirm that BL-918 is a potent activator of ULK1 and autophagy but unable to truly induce mitophagy, therefore, we decided to proceed with this compound in the design of our ULKRECs. For the targeting ligand, we use a 2-phenylindole derivative TSPO ligand, which is known to bind to the translocator protein (also known as, peripheral benzodiazepine receptor) on the mitochondrial outer membrane^35–37^ and has been implemented in AUTAC design previously in the AUTAC4 chimeric molecule^16^.

**Figure 1.**
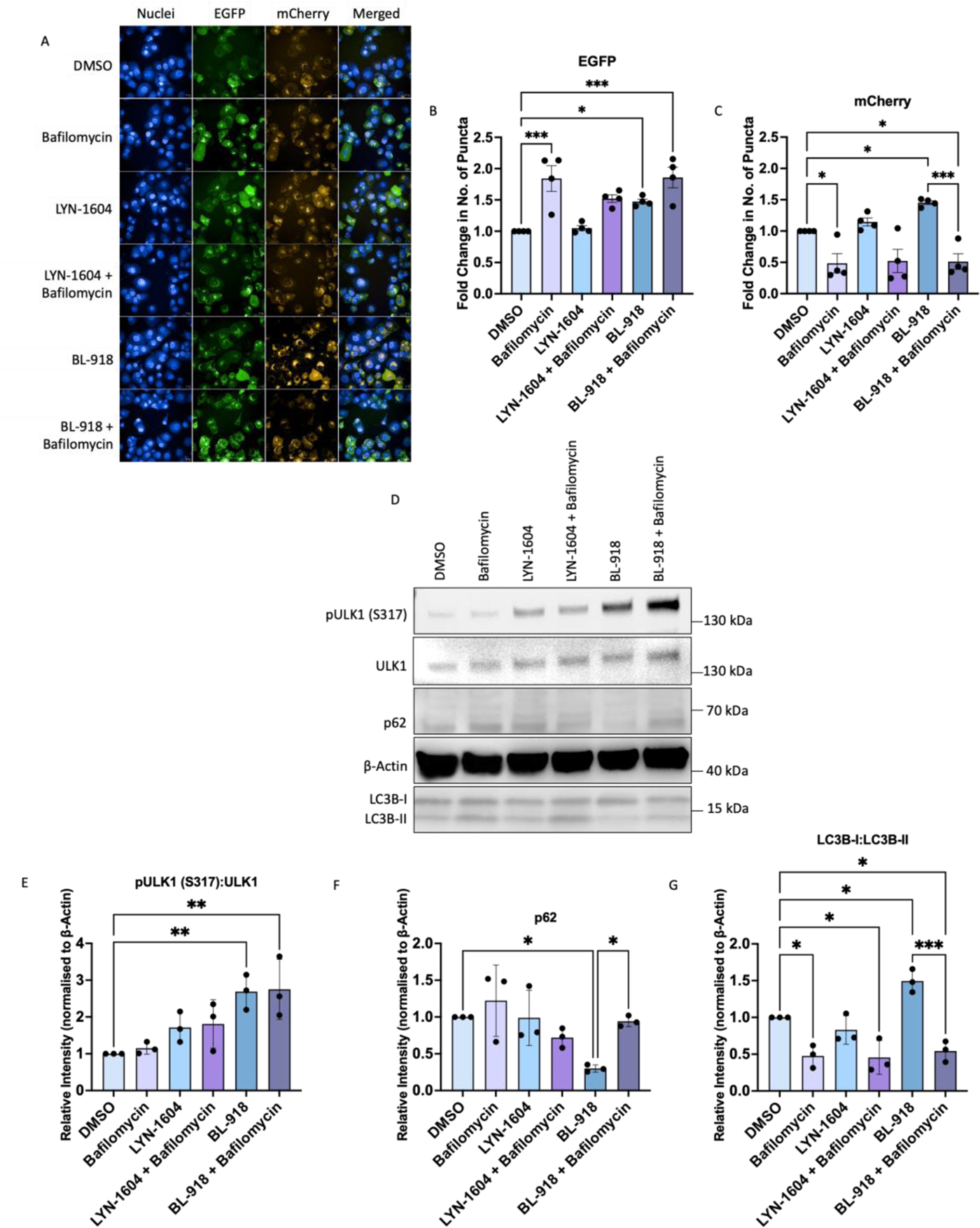
BL-918 is a potent ULK1 agonist and autophagy inducer (A) Representative images of PANC-1 cells stably expressing the mCherry-EGFP-LC3 tandem reporter treated with ULK1 agonists. Nuclei stained with Hoechst 33342, autophagosomes are shown in the EGFP channel and autolysosomes in the mCherry channel. Fold change in the number of autophagosomes (B) and autolysosomes (C) puncta in ULK1 agonist treated cells. (D) Western blot of PANC-1 cells treated with DMSO, bafilomycin and ULK1 agonists ± bafilomycin; probed for p-ULK1 (S317), ULK1, p62, β-actin, LC3-I and LC3-II. Quantifications of western blot for changes in the ratio of p-ULK1 (S317):ULK1 (E), p62 (F) and LC3I:LC3II (G). All data shown are mean ± SD, and at least n = 3, *p<0.05, **p<0.01, ***p<0.001, One-way ANOVA with Šidák multiple comparisons test.

Next, we designed a chimeric compound composed of a BL-918 derivative as our autophagy activator linked to the TSPO ligand as our targeting ligand to the outer mitochondrial membrane, utilising Huisgen–Sharpless click chemistry for the key linkage step^38–41^ . We designed a chemical synthesis route in 9 steps to produce two ULKRECs, NZ-65 and NZ-66. NZ-65 employs a polyethylene glycol (PEG) linker while NZ-66 uses a six-carbon hexane linker (Figure 2A). The resulting molecules were >99% pure as verified by LCMS (see Supplementary Information). Computational modelling shown in Figure 2B suggests that the interactions between BL-918 and ULK1 are maintained with our ULKRECs and ULK1 except for the π-sulfur interaction involving the thiourea in the parent molecule, as our compounds replace this with a urea functional group instead. Importantly, key interactions shown to be necessary for the activation of ULK1, namely, Arg18, Lys50, Asn86 and Tyr89^8^, appear to be maintained, as shown below in the two-dimensional protein-ligand interaction diagram in Figure 2C. When measuring the activation of ULK1 by NZ-65 and NZ-66 in an *in vitro* kinase assay measuring ATP consumption, we find comparable activity of both molecules versus BL-918 (Supplementary Figure 2). Likewise, computational modelling also suggests typical interactions between the TSPO ligand and TSPO also remain unchanged^42,43^ (Figure 2B).

**Figure 2.**
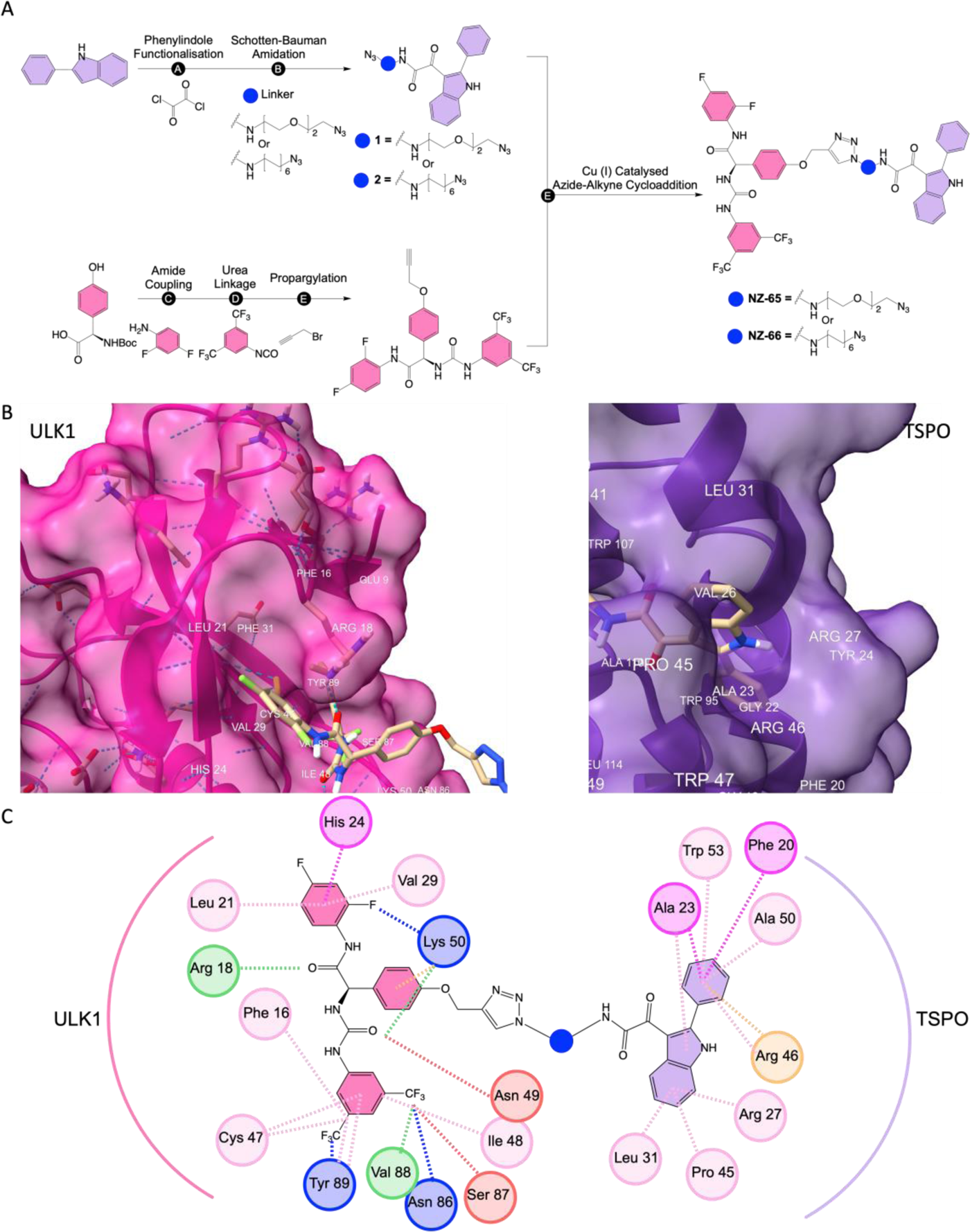
(A) Synthetic route for the synthesis of ULKRECs, **NZ-65** and **NZ-66**, see supporting information for complete synthetic procedure: (**A**) Oxalyl chloride, Et_2_O, RT, 90% (**B**) Azido-PEG2-Amine or 6-Azido-hexylamine, NaHCO_3_, DCM, RT, **1** = 57% and **2** = 67%, (**C**) i. *N-*Methylmorpholine, isobutyl chloroformate, THF, 2,4-difluoroaniline, -20°C to RT.; ii. 4M HCl in 1,4-dioxane = 1/1, RT, 16% (2 steps); (**D**) i. 3,5-bistrifluoromethylphenyl isocyanate, DIPEA, MeCN, 50°C; ii. 2M NaOH/THF = 1/1, RT, 17% (2 steps) (**E**) Propargyl bromide, K_2_CO_3_, MeCN, 80°C; (**F**) Compound **1** or **2**, Compound **3**, THF in H_2_O = 10:1, CuSO_4_, Sodium Ascorbate, 50°C, **NZ-65** = 14%, 2 steps, **NZ-66** = 16%, 2 steps. (B). 3D docking of ULKRECs to ULK1 (Left) and TSPO (Right), poses based upon the kinase domain structure of ULK1 (PDB code: <u>4WNO</u>) and TSPO (PDB Code: 2MGY). (C) Two-dimensional representation of the modelled 3D structures of ULKRECs binding to ULK1 and TSPO proteins, analysed with BIOVIA Discovery Studio. Interactions are shown by dashed lines: Blue = Halogen Interaction/H-bond, Green = Conventional H-bond, Red = C-H bond, Pink = π-alkyl, Magenta = π-π T-shaped or π-Amide stacked, Orange = π-cation

We first investigated whether modifying the ULK1 agonist by the inclusion of a linker and targeting ligand retains agonist activity. When assessing phosphorylation of ULK1 substrates; the auto-phosphorylation of ULK1 at Ser317, we observed a strong increase in phospho-Ser317 levels, suggesting that both NZ-65 and NZ-66 retained the ability to activate ULK1 (Figure 3A-B). In addition to this, we observed increases in the LC3I:LC3II ratio coupled with significant reductions in p62/SQSTM1 (Figure 3C, D), indicative of an induction of autophagy. Using the mCherry-EGFP-LC3 tandem reporter we support these findings, as we describe statistically significant changes, ranging from 1.5 and 1.3 fold increases in the number of EGFP and mCherry puncta, respectively (Figure 3E-G). The enhanced number of mCherry puncta were reversed by co-treatment with the specific autophagy inhibitor bafilomycin A1 which blocks the vacuolar-type H+ ATPase proton pump and prevent autophagosome-lysosome fusion, indicating that NZ-65 and NZ-66 do not block autophagic flux.

**Figure 3.**
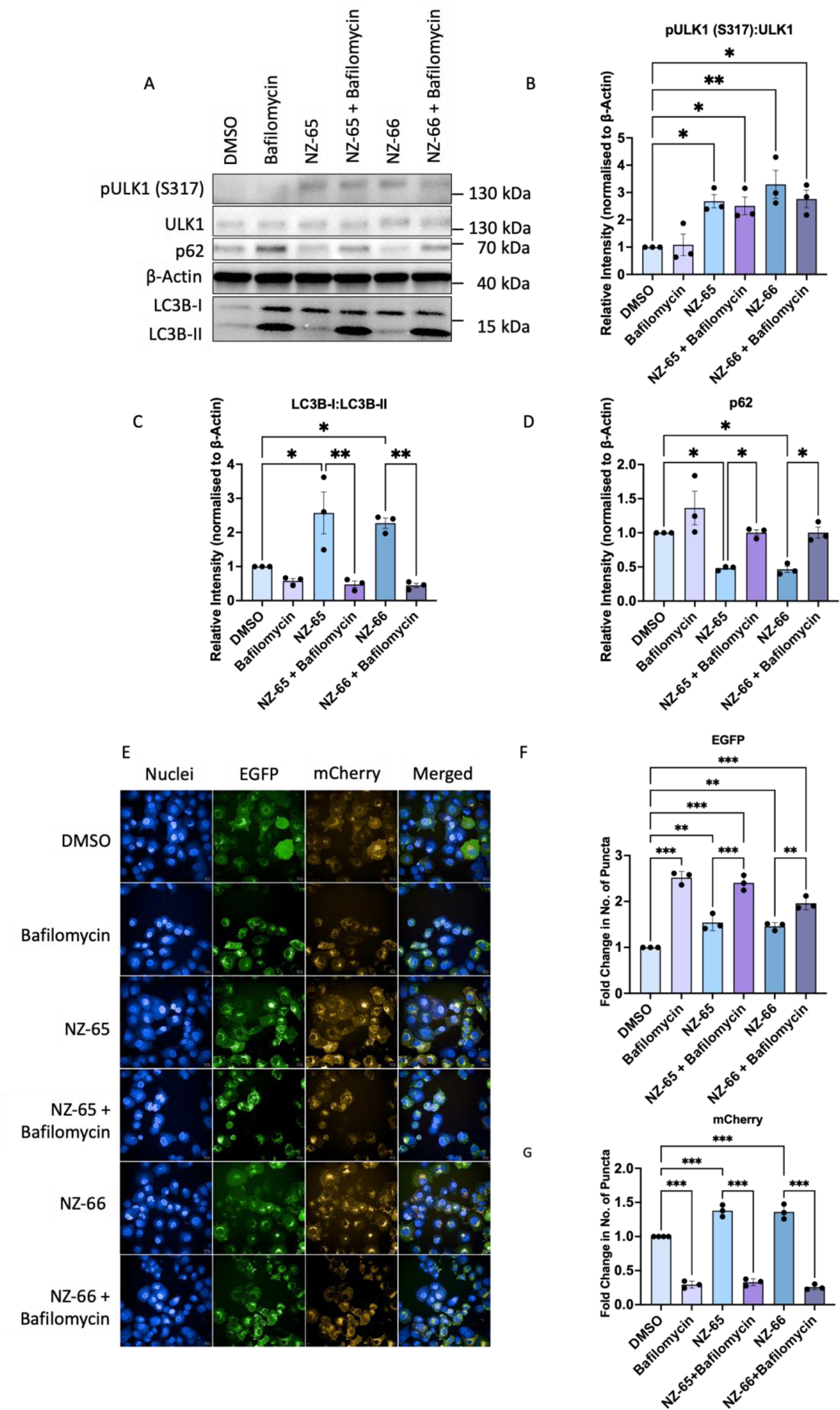
ULKRECs retain ULK1 agonist activity and subsequent induction of autophagy (A) Representative images of PANC-1 cells stably expressing the mCherry-EGFP-LC3 tandem reporter treated with ULKRECs w/ or w/o bafilomycin. Nuclei stained with Hoechst 33342, autophagosomes are shown in the EGFP channel and autolysosomes in the mCherry channel. (B) Fold change in the number of autophagosome (B) and autolysosome (C) puncta. (D) Western blot of PANC-1 cells treated with ULKRECs ± bafilomycin, probed for p-ULK1 (S317), ULK1, p62, β-Actin, LC3I and LC3II. Quantifications of (D) for the pULK1:ULK1 ratio (E) LC3I:LC3II ratio (F) and changes in p62 (G). Data shown are mean ± SD, *p<0.05, **p<0.01, ***p<0.001, One-way ANOVA with Tukey’s multiple comparisons test.

To understand the temporal kinetics of NZ-65 and NZ-66, we assessed the recruitment of ULK1 to mitochondria, using co-localisation of ULK1 to TOM20 by immunofluorescence. We observed an increased co-localisation of ULK1 and TOM20 upon treatment with NZ-65 and NZ-66 after 18 hours of treatment in PANC-1 cells (Figure 4A, B), enhanced by CCCP co-treatment. We also showed, that neither BL-918 nor the TSPO ligand alone induce co-localisation, and cotreatments of ULKRECs with excess TSPO ligand prevents ULK1 co-localisation with TOM20. This suggests that free TSPO receptor is required for ULKREC recognition of mitochondria, and that the observed redirection of ULK1 and colocalisation is facilitated only by the TSPO ligand linked to the ULK1 agonist (Figure 5B). A similar increase in co-localisation was observed in SH-SY5Y cells (Supplementary Figure 3). After showing the co-localisation of ULK1 to the outer mitochondrial membrane (OMM) after ULKREC treatment, we wanted to determine whether the ULKRECs induced mitophagy. To do this, we measured changes in Mfn2 under basal conditions, treatment with ULKRECs and co-treatment with CCCP (Figure 4C, D). We found that the ULKRECs enable the degradation of Mfn2, enhanced upon mitochondrial insult, and additionally, we found that ULKREC-mediated degradation of Mfn2 (and hence, mitophagy) is dependent on autolysosomal activity, as co-treatment with bafilomycin abolishes this activity (Supplementary Figure 4). Similarly, we used the mito-mKeima assay to visualise these changes in mitophagy over time and observed remarkably strong increases in mitophagy upon co-treatment of NZ-65 or NZ-66 with the mitochondrial toxins CCCP and antimycin/oligomycin (Figure 4E-G). Importantly, co-treating with excess TSPO ligand blocks the redirection of ULK1 as previously mentioned, and subsequently restricts the enhancement of mitophagy by either NZ-65 or NZ-66 (Figure 4H, I), again demonstrating targeting-ligand mandated redirection of autophagy. When assessing the mitochondrial membrane potential with TMRM, neither NZ-65 nor NZ-66 showed any effect (Figure 4J), hence the induction of mitophagy is not an artefact of mitochondrial membrane depolarisation. As previously mentioned, it is important to note, that these findings are absent when investigating the mitophagic potential of BL-918; instead, we observe induction of mitophagy only at higher concentrations likely due to loss of mitochondrial membrane potential as seen by a depolarisation of the outer mitochondrial membrane at higher concentrations of BL-918 (see Supplementary Figure 1).

To investigate the sequence of events further we transfected PANC-1 cells with the pMRX-IP-Venus-mULK1 plasmid^31^ for 48 hours before treating cells with ULKRECs. We find that ULK1 recruitment starts approximately 3 hours after treatment and continued to be visible for 18 hours (See Supplementary Video 1 and Supplementary Figure 5). Overall, these results suggest that NZ-65 and NZ-66 work as designed, by recruiting ULK1 to the mitochondria, resulting in a local activation of ULK1 at the mitochondrial membrane and subsequently inducing mitophagy. A mitochondrial insult with CCCP or antimycin/oligomycin further drives the activity of both ULKRECs and enhances mitophagic activity.

**Figure 4.**
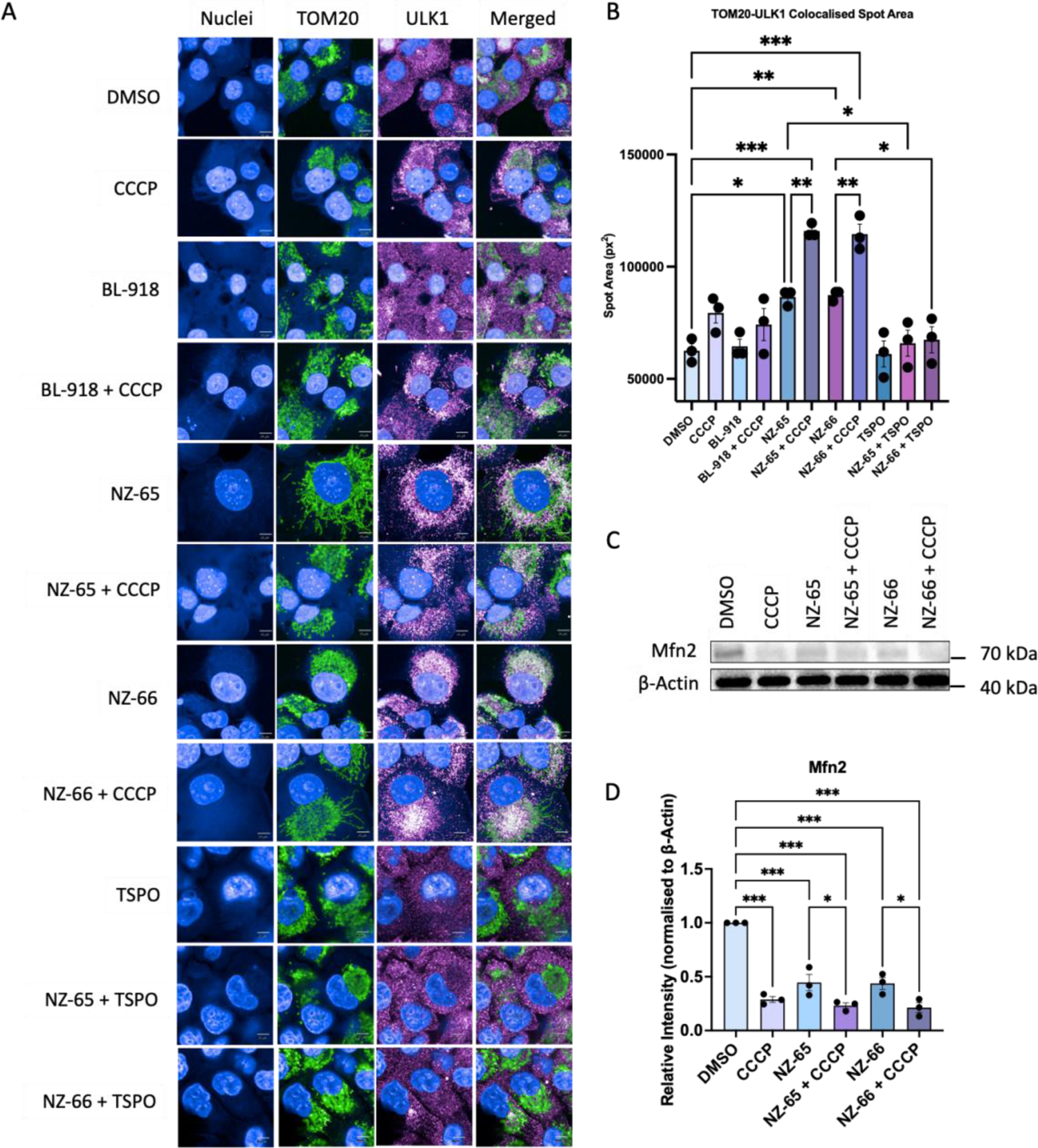
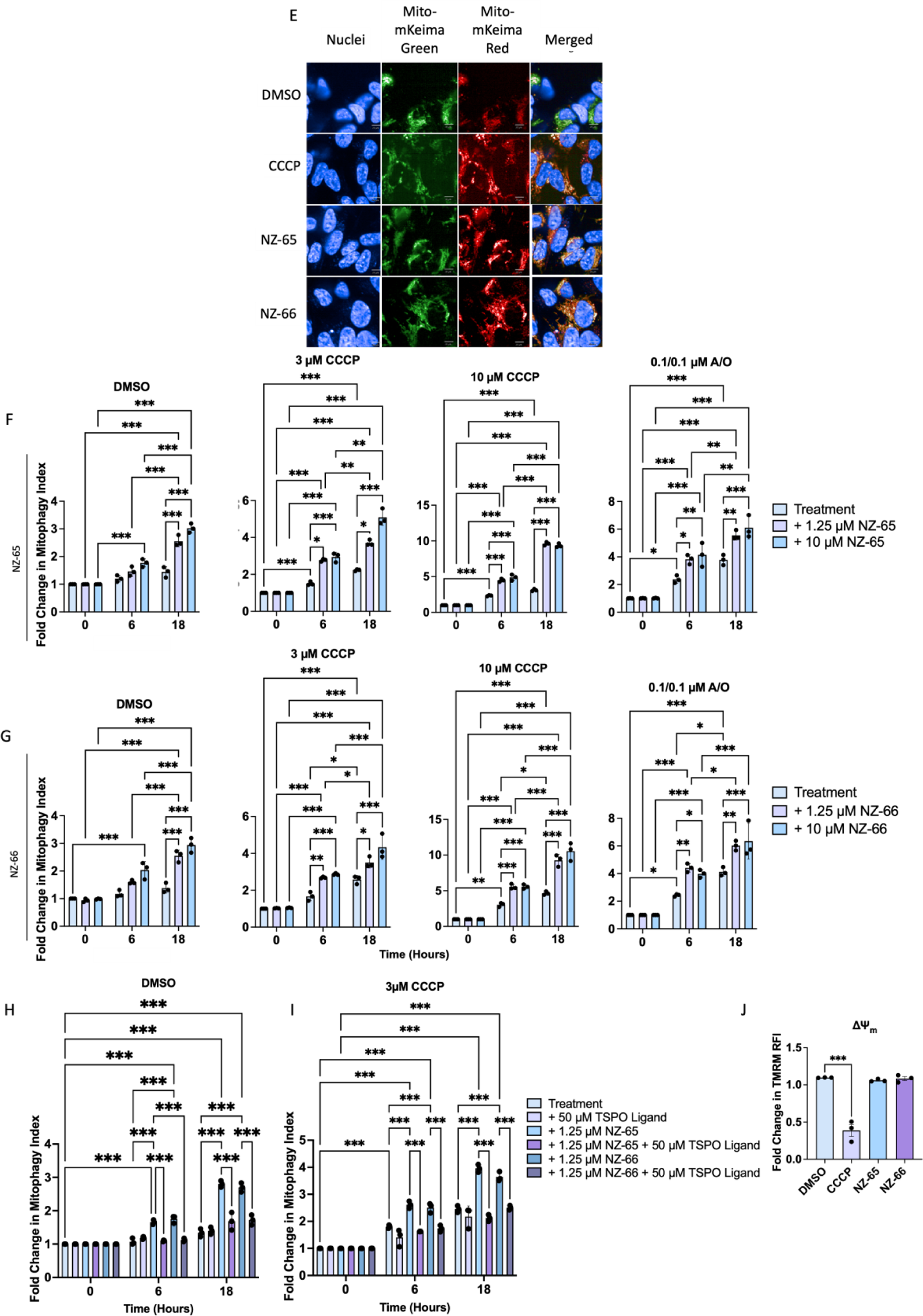
ULKRECs induce potent mitophagy after colocalisation and local activation of ULK1 at mitochondria (A) Representative immunofluorescence of PANC-1 cells treated with CCCP, ULKRECs or both and immunostained for TOM20 and ULK1 and counterstained with Alexa 488 and 568, respectively. Nuclei stained with Hoechst 33342. (B) Quantification of the colocalised ULK1 to TOM20 spot area of PANC-1 cells treated with DMSO, 5 μM CCCP, 10 μM BL-918, 10 μM BL-918 + 5 μM CCCP, 1 μM NZ-65, 1 μM NZ-65 + 5 μM CCCP, 1 μM NZ-66, 1 μM NZ-66 + 5 μM CCCP, 50 μM TSPO ligand, 1 μM NZ-65 + 50 μM TSPO ligand, 1 μM NZ-66 + 50 μM TSPO ligand. (C) Western blot of SH-SY5Y cells treated with 1 μM ULKRECs, 5 μM CCCP or both, probed for Mfn2, β-Actin and Tom20 – quantifications are shown in (D) for Mfn2, normalised to Actin, n = 5. (E) Representative images of SH-SY5Y cells stably expressing the mito-mKeima reporter treated with 3 μM CCCP, NZ-65 or both. Nuclei were stained with Hoechst 33342, healthy mitochondria are shown in green and mitochondria in lysosomes are shown in red. Fold change in mitophagy index over time after treatment for the indicated timepoints with title treatments alone or co-treated with 1.25 μM ULKREC or 10 μM ULKREC NZ-65 (F) or NZ-66 (G). Fold change in mitophagy index over time after treatment for the indicated timepoints with DMSO, 1.25 µM NZ-65 or NZ-66 ± 50 µM TSPO ligand under co-treatment with DMSO (H) or 3 µM CCCP (I). (J) Fold change in TMRM relative fluorescent intensity (and hence ΔΨ_m_) after 1h treatment with DMSO, 3 μM CCCP or 20 μM ULKREC. Normalised to number of nuclei. All data shown are mean ± SD, n = 3, *p<0.05, **p<0.01, ***p<0.001. For colocalisation experiments one-way ANOVA with Šidák multiple comparisons test. For TMRM, One-way ANOVA with Dunnett’s multiple comparisons test. For western blots, one-way ANOVA with Tukey’s multiple comparisons test. For mito-mKeima experiments, Two-way ANOVA with Tukey’s multiple comparisons test.

To confirm whether the activity of ULK1 is required for NZ-65 and NZ-66, we studied the recruitment of LC3 to mitochondria in ULK1/2^-/-^ mouse embryonic fibroblasts^27^ (gift from Professor Sharon Tooze, Francis Crick Institute) and the degradation of Mfn2. In accordance with previous results, we observed degradation of Mfn2 upon treatment with NZ-65 and NZ-66 that was further enhanced by co-treatment with CCCP; in contrast, we observed no degradation of Mfn2 in the ULK1/2^-/-^ double knockout cells upon ULKREC treatment (Figure 5A, B). We also observed an increase in co-localisation of LC3 to the mitochondrial marker ATP5a after treatment with NZ-65 and NZ-66, that was absent in the ULK1/2^-/-^ cells (Figure 5C-E), suggesting recruitment of the LC3 cargo protein to the mitochondria when MEFs are treated with NZ-65 or NZ-66, and that this is reliant on the recruitment and activation of ULK1 at the OMM. These findings were cross-validated with the recently developed dual ULK1/2 inhibitor, SBP-7455^6^, additionally demonstrating the requirement for ULK1 and canonical autophagy signalling for ULKREC mediated degradation of Mfn2 and hence, mitochondria (Supplementary Figure 6). Therefore, we conclude that NZ-65 and NZ-66 are novel ULKREC molecules that initiate the recruitment of ULK1 to mitochondria, leading to an increase in local ULK1 activity, the local formation of an autophagosome and mitophagy and, that their activity is inherently reliant on autophagy. Interestingly, it appears that ULK1/2 activity is required for toxin-induced mitophagy, and furthermore, NZ-65/66 enhance basal mitophagy though further enhance it in response to insult to prepare the mitochondria for uptake in an autophagosomes.

**Figure 5.**
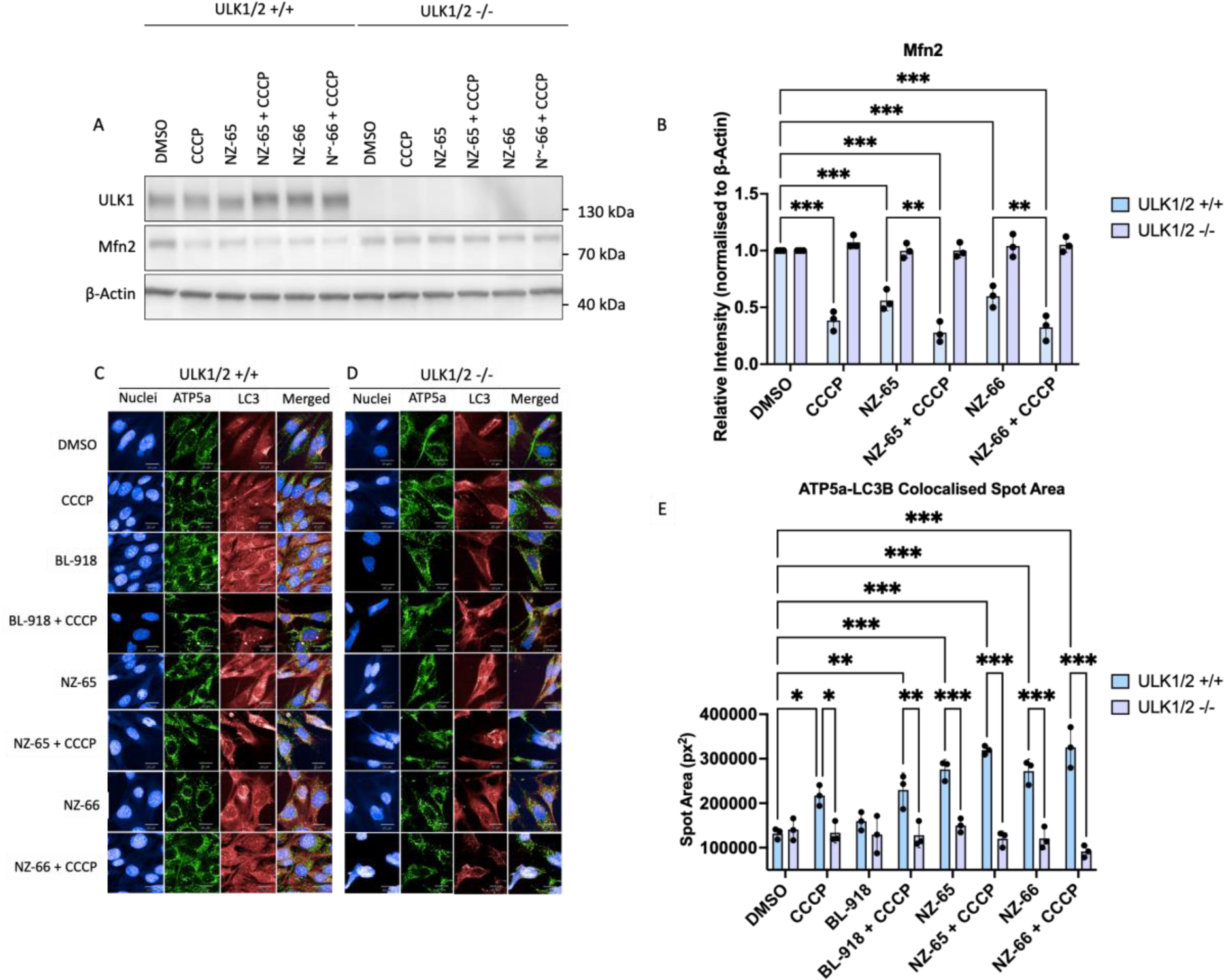
ULKRECs rely on ULK1 for the redirection of autophagy components to mitochondria and subsequent mitophagic degradation (A) Western blot of mouse embryonic fibroblasts (MEF) WT and MEF ULK1/2^-/-^ treated with 1 μM ULKRECs, 5 μM CCCP or both and probed for ULK1, Mfn2 and Actin – quantification of the relative intensity of Mfn2 shown in (B), normalised to Actin. Representative immunofluorescence of MEF WT (C) and MEF ULK1/2^-/-^ (D) treated with 5 μM CCCP, 1 μM ULKRECs or both and immunostained for Alexa 488 ATP5a and Alexa 647 LC3B. Quantification of the colocalised LC3 to ATP5a spot area shown in WT vs ULK1/2^-/-^ treated MEF cells shown in (E). Data shown are mean ± SD, n = 3, *p<0.05, **p<0.01, ***p<0.001, Two-way ANOVA with Tukey’s multiple comparisons test.

To explore the potential therapeutic benefit of novel chimeric molecules, enabling the potent exogenous induction of mitophagy, we investigated whether the ULKRECs were able to induce mitophagy in Parkinson’s disease (PD) patient-derived PINK null fibroblasts. A loss of function over time and/or mutations in PINK/PRKN have been associated with PD and can predispose individuals to hereditary PD^44–48^. Treatment with FCCP in WT fibroblasts caused a substantial increase in pUb (S65) levels, which did not occur in the PINK-null cells (Figure 6A) – this was expected as the PINK-null cells are unable to signal for mitophagy in response to stressors. We also found that treatment with ULKRECs does not activate phosphorylation of ubiquitin at S65, suggesting they do not activate PINK1. However, we continued to observe significant reductions in Mfn2, suggesting that the ULKRECs induce mitophagy independent of the PRKN/PINK signalling axis (Figure 6B, C). In agreement with previous experiments, we investigated the colocalisation of ULK1 to TOM20 in both WT and PINK-null cells and correlate the enhanced colocalsation of ULK1 to TOM20 to the observed Mfn2 degradation in WT and PINK-null cells. In PINK-null cells, unable to signal for canonical mitophagy we do not observe colocalisation of ULK to the mitochondria validating this assay (Figure 6D, E)

**Figure 6.**
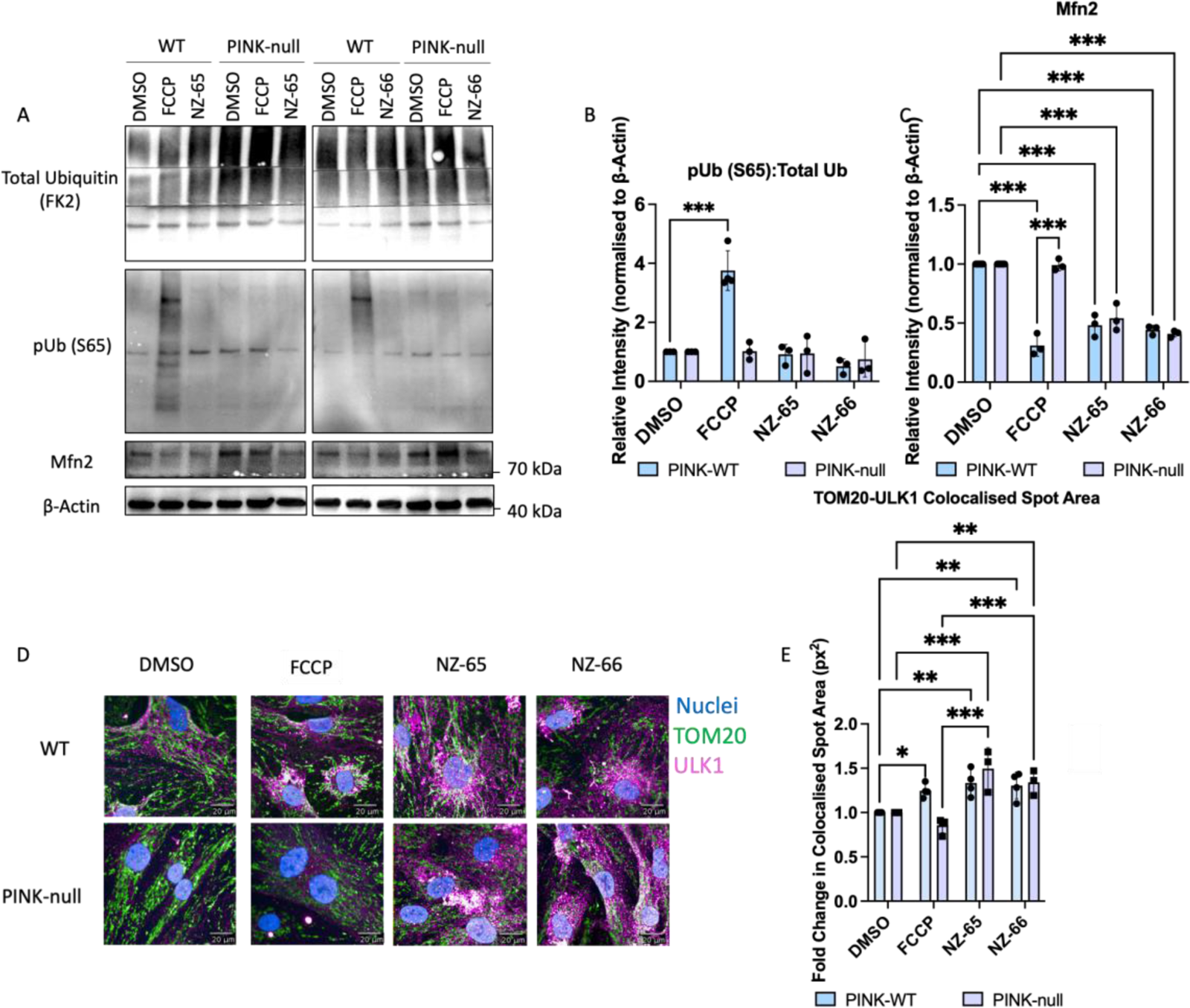
ULKRECs restore mitophagic activity in PINK-null PD patient fibroblasts (A) Western blot of patient derived fibroblasts, healthy control vs. PINK null, treated with 1 μM NZ-65 or NZ-66 for 18h or 10 µM FCCP for 2h and probed for total Ub, pUb(S65), Mfn2 and β-Actin with quantification for the changes in the pUB (S65):Total Ub ratio shown in (B) and Mfn2 (C), respectively normalised to β-Actin. (D) Representative immunofluorescence of primary human fibroblasts isolated from wild type controls and PINK-null Parkinson’s disease patients, treated with 10 µM FCCP, 1 µM NZ-65 or 1 µM NZ-66 and immunostained for TOM20 and ULK1; counterstained with Alexa 488 and Alexa 568 antibodies. Nuclei stained with Hoechst 33342. The quantification of these immunofluorescence experiments is shown in (E). Data shown are mean ± SD, n = 3, *p<0.05, **p<0.01, ***p<0.001, Two-way ANOVA with Tukey’s multiple comparisons test.

## DISCUSSION

In this study, we developed a novel strategy for autophagy-mediated degradation of a cargo as an alternative to PROTAC and AUTAC design with a tractable and clearly described mechanism of action. While PROTACs are suitable for the degradation of proteins and peptides, they cannot be employed for the degradation of non-protein molecules, larger complexes, insoluble proteins aggregates or organelles. Therefore, the ability of the autophagosome to deliver large cargo with a wider substrate scope to the lysosome for degradation can be harnessed through the development of AUTACs. Current strategies using autophagy for targeted degradation generally bind a cargo protein and tether the target of interest (TOI) to key proteins of a forming autophagosome such as LC3^15,17,18^, or p62^19^ for capture by a forming autophagosome for degradation after fusion with lysosomes. Indeed, this tether strategy has demonstrated autophagy can be exploited for the degradation of lipid droplets^18^, protein aggregates and small proteins^17,19^. Although it is expected that tethers would be able to degrade all other autophagy substrates including organelles such as mitochondria, this has not yet been shown. One concern is that this strategy makes use of basal autophagy and the effects on the function of homeostatic autophagy and competition with native autophagy cargo are not known. An alternative strategy developed by Takahashi et al.,^16^ mimics the selective S-guanylation of bacterial envelope proteins during xenophagy^49^ for autophagic degradation to develop AUTAC4, a chimeric molecule consisting of a pseudo-guanylation tag linked to a translocator protein (TSPO) ligand to mark mitochondria for targeted degradation. Although this study demonstrates the usability of the TSPO ligand to target the outer mitochondrial membrane, the mechanism of action of these AUTACs is very poorly understood and demonstrate limited potency in the original article. Because the ULKRECs use the same targeting ligand as AUTAC4, we find that ULKRECs are a superior AUTAC strategy and more potent degradation is achieved when creating autophagosomes at the target site as opposed to marking the TOI for degradation, as potent mitophagy is achieved even at 10-fold lower concentrations^16^.

The current AUTAC strategies have demonstrated the feasibility of delivering cargo (both large organelles and smaller proteins) to the lysosome for degradation, however there are several limitations. First, it is not known how trafficking of the cargo to the autophagosome might impact the efficiency of this process. Secondly, using existing autophagosomes for degradation may be associated with limitations in autophagosome availability, competition with other cargo and possibly saturation of the basal autophagy machinery. We wanted to overcome these issues by developing a novel AUTAC strategy with a tractable and clearly defined mechanism of action. We hypothesized that an autophagosome can be initiated locally by activation of the ULK1 kinase. This is based on previous observations that the activation of ULK1 can trigger a phagophore locally, using a genetic tool based on a chemically induced dimerization assay^26^. The authors demonstrated that ectopic recruitment of NDP52 to mitochondria can trigger mitophagy in an ULK1-dependent manner, mediated by de novo LC3-independent selective autophagy. In this case, ULK1 was able to auto-activate on cargo, an observation that has also been made for ATG1 during cytosol-to vacuole targeting and pexophagy^50,51^. Furthermore, direct targeting of ULK1 to peroxisomes was shown to trigger pexophagy^26^, thus demonstrating that ULK1 localisation can auto-activate the initiation of autophago-lysosomal degradation. The precise mechanism how ULK1 can autoactivate in this context requires further investigation.

Here, we sought to make use of this observation by developing ULKRECs, specific chimeric molecules that link ULK1 and the cargo chemically as a novel targeted degradation strategy. While ULK1 binding molecules may be sufficient to achieve this, we reasoned that the use of an ULK1 activator may be even more beneficial. To do this, we evaluated two known ULK1 agonists LYN-1604^7^ and BL-918^8^. One caveat of LYN-1604 is that LYN-1604 induces cytotoxic autophagy in an MDA-MB-231 cell line and xenografts, raising concerns over its suitability as a therapeutic candidate in diseases where cell death is not desired^7^. BL-918, on the other hand, is an inducer of ULK1 and autophagy that exerts cytoprotective effects in a model of Parkinson’s disease and amyotrophic lateral sclerosis^8,52^. The ULKRECs, NZ-65 and NZ-66 were derived from BL-918: the original compound was modified in the synthesis of our ULKRECs, while comparably retaining its ULK1 binding and activation potential (see Supplementary Figure 2 for in vitro ULK1 activation comparison). We replaced the thiourea functional group with the urea counterpart and at the central free aromatic ring, we converted this into a prop-2-yn-1-yloxyl benzene for the subsequent azide-alkyne Huisgen-Sharpless dipolar cycloaddition. We decided that this was the most appropriate location to attach the linker as computational modelling of BL-918 by Ouyang et al.^8^ suggests that this aromatic ring does not interact with ULK1. We expect that further modifications of our derivative or newer ULK1 agonists will improve the properties of NZ-65 and NZ-66 in the future, making ULKRECs possibly amenable to *in vivo* applications.

BL-918 has been comprehensively studied *in vitro* and *in vivo* for the induction of cytoprotective autophagy in Parkinson’s disease (PD)^8^, and more recently, amyotrophic lateral sclerosis (ALS)^52^. Although BL-918 has been described to protect against neurotoxin-induced parkinsonism, no studies have reported its effect on genetic models of the disease, for example, in PINK-null cells. It is unlikely that a general upregulation of autophagy is sufficient to drive a sole sufficient enhancement of mitophagy, rather enhancing total autophagic activity instead with coincidental mitophagic degradation. Indeed, mito-mKeima experiments using BL-918 demonstrates this (Supplementary Figure 1) while the compound only drives mitophagy after mitochondrial depolarisation at higher concentrations (Supplementary Figure 1). Using the novel ULKRECs NZ-65 and NZ-66, we offer a specific and targeted approach to enhance autophagy after the machinery arrives at the target site, resulting in the desired specific induction of mitophagy as evidenced by the dramatic fold changes in mitophagic activity we report. The current view is, pathological mutations in the PINK/PRKN axis or generally dysfunctional/inefficient mitophagy promotes the accumulation of damaged mitochondria and the aggregation of alpha-synuclein causing subsequent dopaminergic neuronal damage and/or death, resulting in the observed characteristics of the disease^33,44,46–48,53^. Given that coincidental degradation by BL-918 likely contributes to its beneficial cytoprotective effects in the reported disease models, it is then expected that the targeted autophagic degradation of TOIs by ULKRECs would be more effective and/or potent. More specifically, it Is also expected that a targeted induction of mitophagy through ULKRECs would be more therapeutically relevant versus a general inducer of autophagy such as BL-918 in the case of PD; whether targeting one of the many root pathological mechanism of PD by the exogenous induction of mitophagy, achieved using NZ-65 and NZ-66 in mitophagy-deficient PD patient cells yields actual therapeutic benefit, remains to be seen. Similarly, one could argue that the general upregulation of autophagy in the ALS study using BL-918, coincidentally results in the clearance of toxic SOD1 aggregates^52^ and a more targeted approach such as a SOD1 ULKREC, could potentially be more potent and therapeutically relevant. As we demonstrated that the ULKRECs bypass conventional mitophagy signalling, other forms of mitophagy such as FUNDC1-mediated^54^ mitophagy may equally benefit, for example, enhancing mitophagy to protect cardiac function under ischemia or reduce cardiac hypertrophy after pathological cardiac stress^55,56^.

It has been shown that the chemically induced targeting of ULK1 to mitochondria using the FKBP-dimerization system resulted in an activation of PINK/Parkin independent mitophagy^26^. Interestingly, the activity of ULK1 was required for mitophagy in this model. This agrees with our observation that there was no degradation of Mfn2 even upon mitochondrial insult with CCCP in ULK1/2^-/-^ or SBP-7455 treated cells suggesting ULK family proteins are essential for mitophagy. This finding is supported by work demonstrating that FUNDC1-mediated mitophagy does not occur in ULK1/2^-/-^ cells and that this is rescued with ULK1 expressing constructs^57^. These results suggest a more general requirement of ULK1/2 in mitophagy processes and highlights the centrality of ULK1/2 proteins in autophagy processes. Generally, incorporation of fragmented mitochondria in an autophagosome is largely favoured over the capture of whole large components^58–60^. This may explain why we see that mitochondrial damage enhances mitophagy in ULKREC co-treated cells, likely by inducing mitochondrial fragmentation as well as from enhanced autophagic activity that is subsequently redirected by ULKRECs. As mentioned previously, we also observed a significant enhancement of basal mitophagy upon sole ULKREC treatment through the redirection of ULK1 and subsequent activation. We also showed that ULKRECs do not affect the mitochondrial membrane potential, and do not adversely affect cell viability at experimentally relevant concentrations (Supplementary Figure 7).

To conclude, prototypical ULK1 targeted chimeras NZ-65 and NZ-66 redirect and colocalise ULK1 to the outer mitochondrial membrane via the linked translocator protein ligand. Targeted localised activation of ULK1 at the OMM by ULKRECs exogenously induces mitophagy, enhanced further by mitochondrial insult and, as the PINK/PRKN axis is bypassed by this exogenous induction of mitophagy, the ULKRECs enable mitophagic activity in PINK-null mitophagy-deficient Parkinson’s disease patient cells.

## ACKNOWLEDGEMENTS

We would like to thank Eliona Tsefou, Nivedita Singh, Janos Kriston-Vizi, Edith Chan and Katherine Morling for advice and comments. We thank Sharon Tooze for ULK1/2 double knockout mouse embryonic fibroblasts^27^. This work was funded by the Medical Research Council (Ref MC_U12266B, MR/M02492X/1), the UCL Therapeutic Innovation Network initiative with funding from MRC Confidence in Concept 2017 UCL MC/PC/17180, the Biotechnology and Biological Sciences Research Council (BBSRC) (BB/T008709/1) the Wellcome Collaborative award (214344, DLS), Wellcome Trust Rapid Response call 204841/Z/16/Z and UCL MRC Confidence in Concept 2020 UCL MC/PC/17180 award through UCL (Small Molecules) Therapeutic Innovation Network.

## CONTRIBUTIONS

NZ, DLS and RK designed the experiments. NZ performed the biological experiments and chemical synthesis of compounds. NA developed the stable cell line of PANC-1 expressing mCherry-EGFP-LC3. VP and DLS planned the synthetic route and synthesis of compounds. RK and DLS conceived and directed the study. All authors were involved in writing and editing the manuscript.

