## Supplemental Information for "Development of the ULK1-Recruiting Chimeras (ULKRECs) to enable proximity-induced and ULK1-dependent degradation of mitochondria"

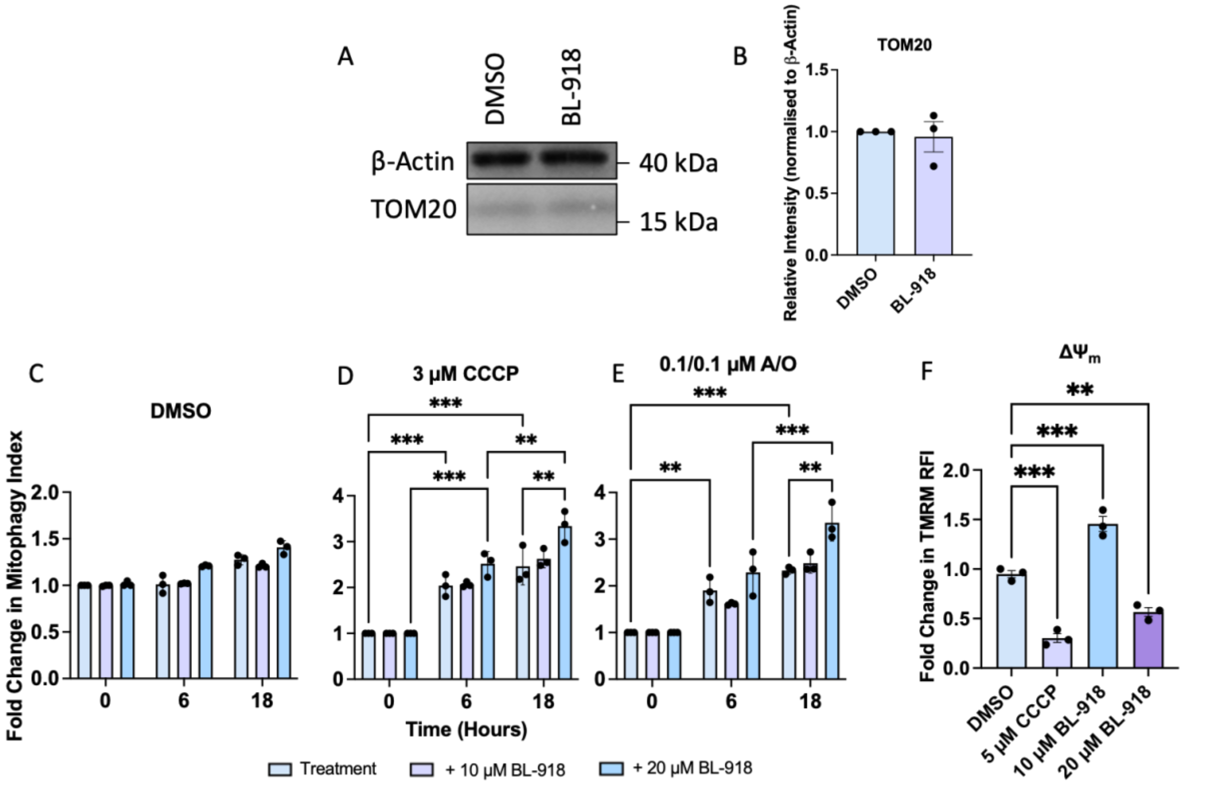


Supplementary Figure 1. BL-918 is unable to induce mitophagy at experimentally relevant concentrations. (A) Western blot of SH-SY5Y cells treated with 10 mM BL-918 and probed for Vinculin and TOM20; quantification shown in (B) was normalised to Vinculin. (C-E) Fold change in mitophagy index over time of SH-SY5Y cells treated with 0 µM, 10 µM and 20 µM BL-918 for the indicated timepoints, co-treated with DMSO, 0.1/0.1 µM A/O or 3 µM CCCP, respectively. (F) TMRM relative fluorescent intensity after 1h treatment with DMSO, 10 µM CCCP, 10 µM and 20 µM BL-918. Normalised to number of nuclei. All data shown are mean ± SEM, n = 3, *p<0.05, **p<0.01, ***p<0.001. For TMRM experiments, one-way ANOVA with Dunnett's multiple comparisons test. For western blots, one-way ANOVA with Tukey's multiple comparisons test and unpaired t-test. For mito-mKeima experiment, two-way ANOVA with Tukey's multiple comparisons test. BL-918 is unable to induce mitophagy at experimentally relevant concentrations – exceeding this, results in mitochondrial depolarisation induced mitophagy and hence an artefact of BL-918 rather than intended activity.

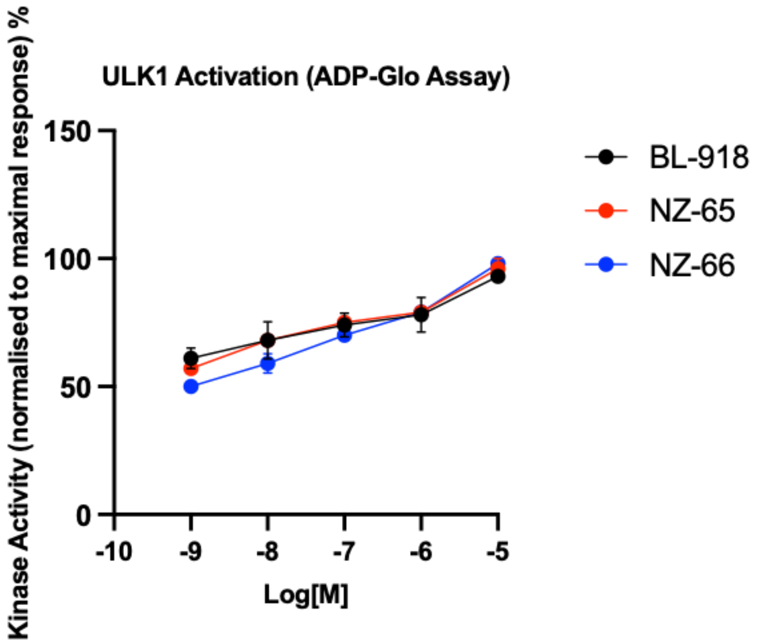


Supplementary Figure 2. ULKRECs retain ULK1 agonist activity, comparable to BL-918. ULK1 kinase activity was measured using the ADP-Glo assay after 1h treatments with increasing concentrations of BL-918, NZ-65 or NZ-66. Relative kinase activity was normalised to the maximal response. Data shown are mean ± SEM, n=3, one-way ANOVA, Dunnett's multiple comparisons test.

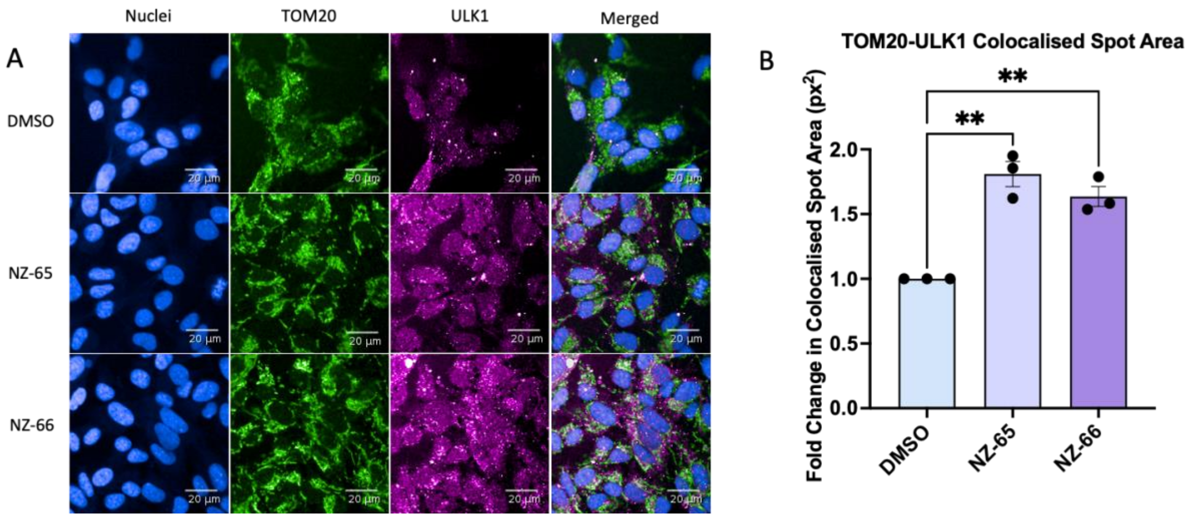


Supplementary Figure 3. ULKRECs induce colocalisation of ULK1 to the outer mitochondrial membrane in SH-SY5Y cells. Representative immunofluorescence of SH-SY5Y cells treated with DMSO, 1 µM NZ-65 or 1 µM NZ-66 and immunostained for TOM20 and ULK1; counterstained with Alexa 488 and 568 respectively. Nuclei stained with Hoechst 33342. (B) Quantification of the colocalised ULK1 to TOM20 spot area in SH-SY5Y cells with treated with the previously described. Data shown are mean ± SEM, n=3, two-way ANOVA, Tukey's multiple comparisons test. **p<0.01. NZ-65 and NZ-66 induce significant co-localisation of ULK1 to TOM20 – no significant difference between NZ-65 and NZ-66.

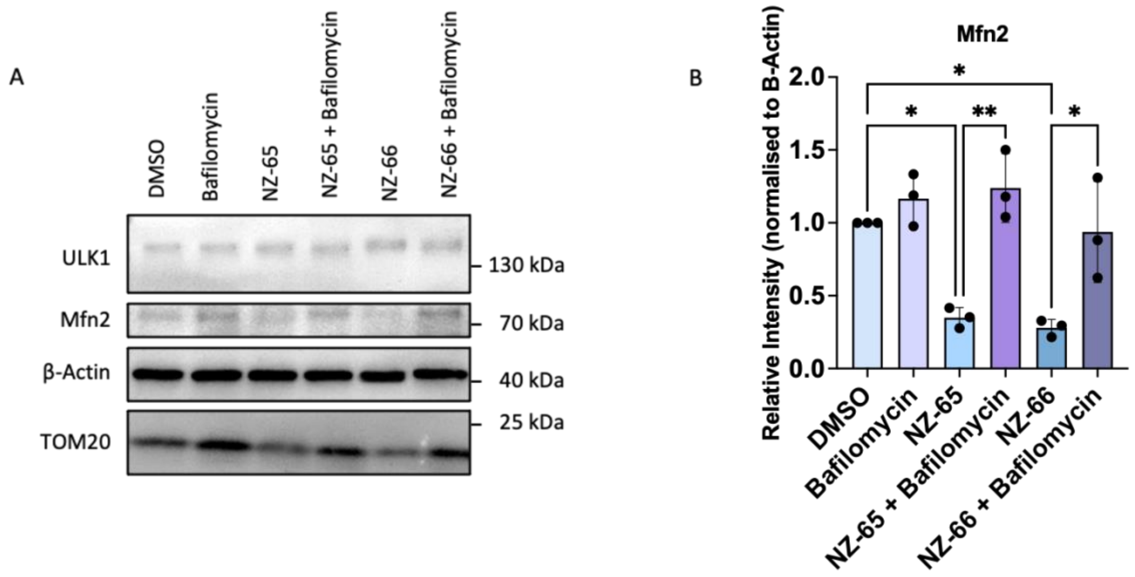


Supplementary Figure 4. Mitochondrial degradation by ULKRECs is lysosomally dependent and inhibited by bafilomycin. (A) Western blot of PANC-1 cells treated with DMSO, 10 nM bafilomycin, 1 μM NZ-65/NZ-66 ± 10 nM bafilomycin, probed for ULK1, Mfn2, β-actin and TOM20. Quantification of western blot for changes in Mfn2 shown in (B). Data shown are mean ± SEM, n = 3, two-way ANOVA, Tukey's multiple comparisons test. *p<0.05, **p<0.01. NZ-65 and NZ-66 induce dramatic changes in Mfn2 levels, suggestive of mitophagy; this is abolished on co-treatment with bafilomycin, demonstrating autolysosomal requirement for the mechanism of action.

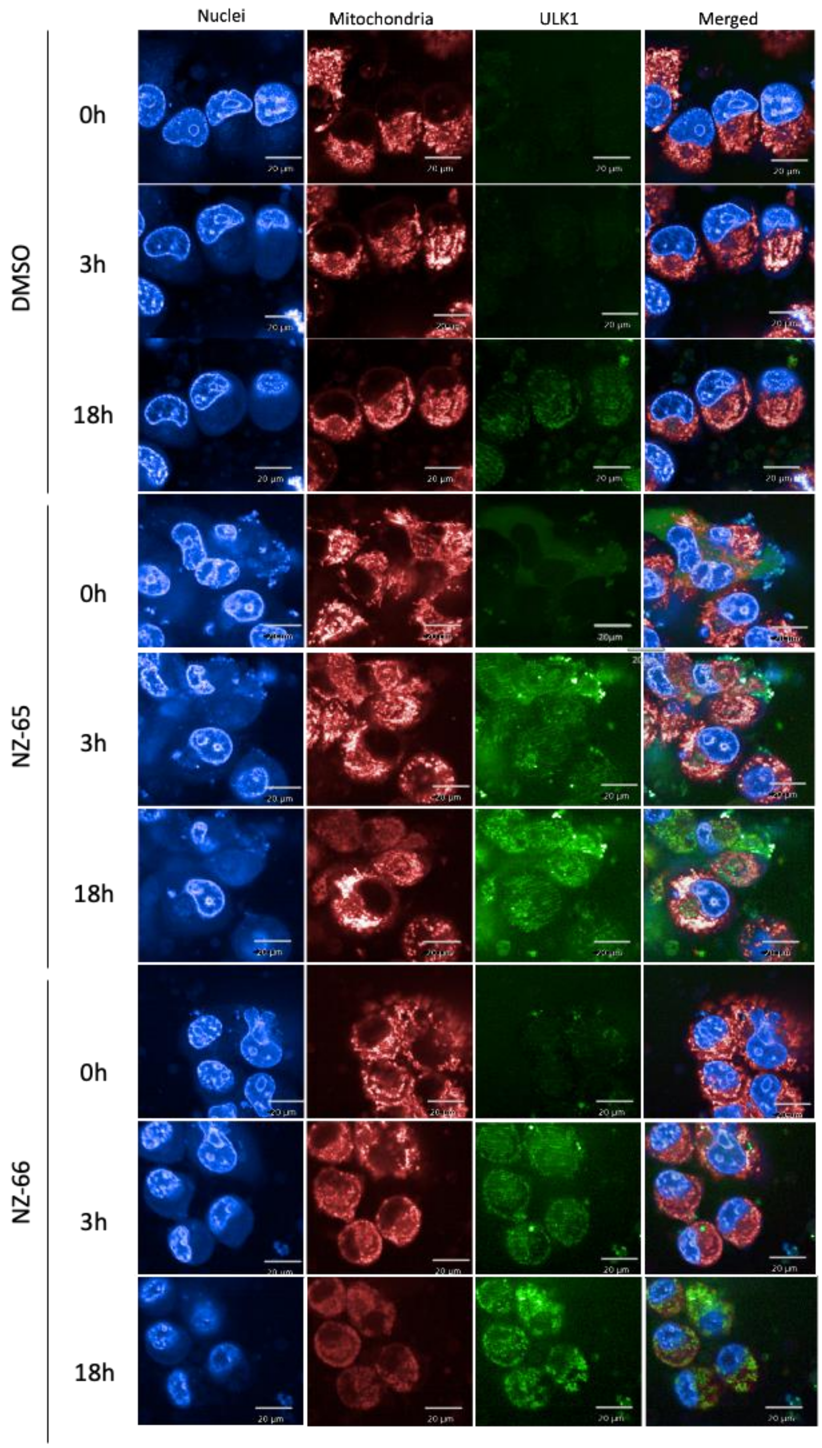


Supplementary Figure 5. ULKREC-mediated colocalisation of ULK1 to mitochondria begins approximately 3 hours after treatment, persisting for at least 18 hours post-treatment. PANC-1 cells were transfected pMRX-IP/Venus-mULK1 with lipofectamine 3000 in antibiotic-free Opti-MEM media for 48 hours prior to treatment and subsequent image capture at hourly intervals. Representative images are shown at 0h, 3h and 18h after treatment.

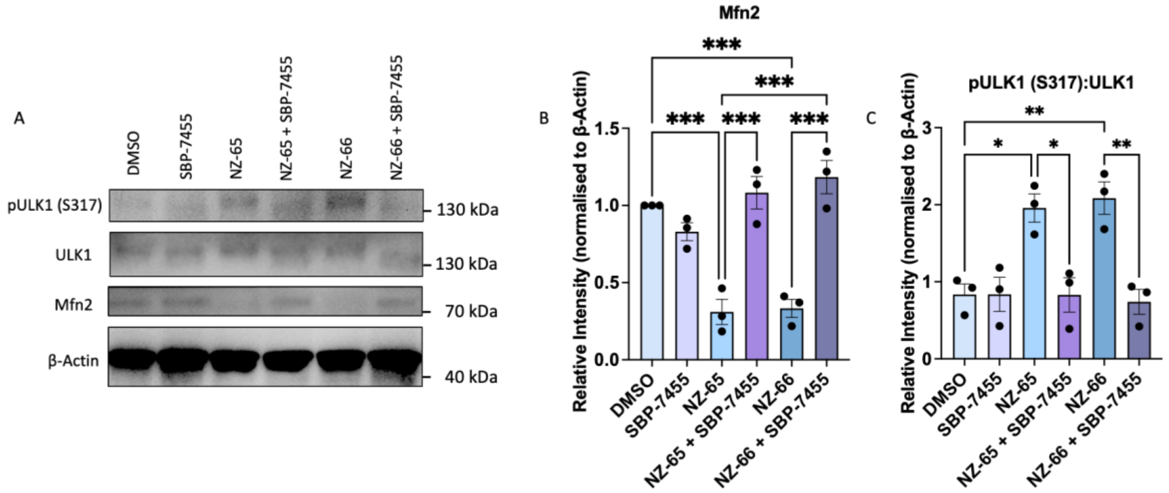


Supplementary Figure 6. ULKRECs require ULK1 for activity. (A) Western blot of PANC-1 cells treated with 1 μM ULKRECs, 5 μM CCCP or both in the presence or absence of the dual ULK1/2 inhibitor SBP-7455 (5 μM) and probed for pULK1(S317), ULK1, Mfn2 and Actin – quantifications shown in (B) and (C) for changes in Mfn2 andpULK1 (S317):ULK1 ratio after treatments, respectively, normalised to Actin. Data shown are mean ± SEM, n = 3, *p<0.05, **p<0.01, ***p<0.001. One-way ANOVA with Tukey's multiple comparisons test.

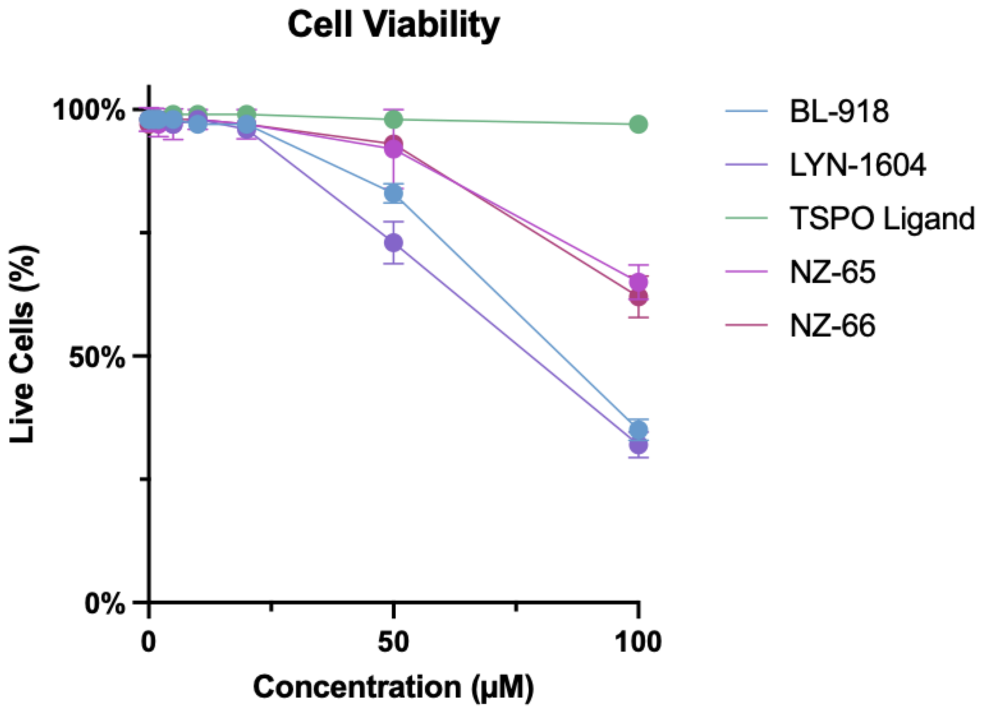


Supplementary Figure 7. Cell Viability of PANC-1 cells treated with increasing concentrations of BL-918, LYN-1604, TSPO Ligand, NZ-65 or NZ-66. Cell viability was determined using a DRAQ7 assay and 100% refers to no observable DRAQ7 signal with Mean RFI greater than background.

***N*-(2-(2-(2-azidoethoxy)ethoxy)ethyl)-2-oxo-2-(2-phenyl-1H-indol-3-yl)acetamide**

**1H NMR**

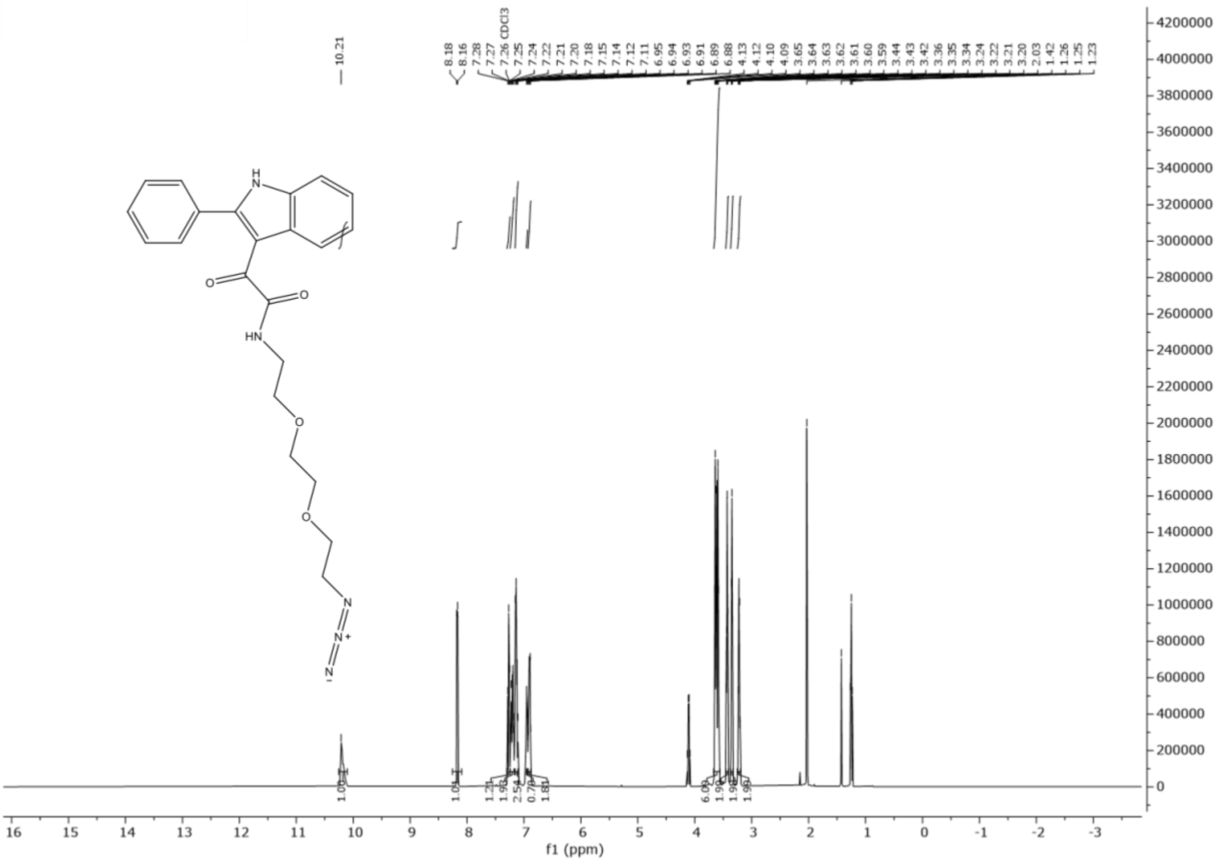

$^{1}$H-NMR (500MHz, MeOD, *J* Hz) $\delta$ ppm 8.99 (s, 1H), 8.18 (d, 1H, *J* = 7.70 ), 7.42 (q, 3H, *J* = 6.44 ), 7.30 (quintet, 5H *J* = 7.29), 6.80 (t, 1H, *J* = 5.75), 3.25 (t, 2H, *J* = 6.90), 3.11 (q, 2H, *J* = 6.77), 1.57 (quintet, 2H, *J* = 7.15), 1.46 (t, 2H, *J* = 7.33), 1.39 – 1.21 (m, 4H).

**13C NMR**

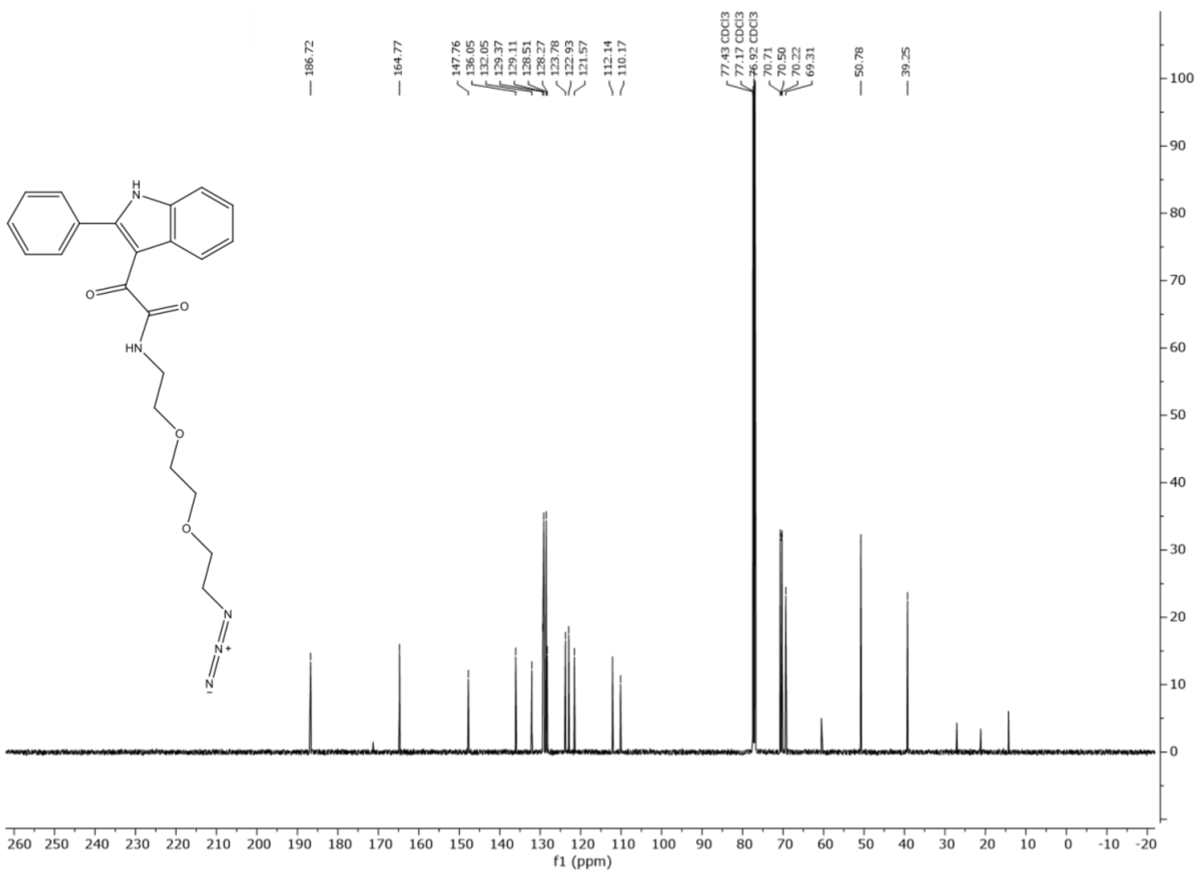


$^{13}$C-NMR (500MHz, MeOD) δ ppm 186.87, 163.81, 147.44, 135.69, 132.46, 129.64, 129.08, 128.71, 128.21, 124.09, 123.10, 121.98, 111.37, 110.64, 51.48, 39.37, 29.81, 29.20, 28.84, 26.53, 26.49.

**UV-Chromatogram (LC-MS)**

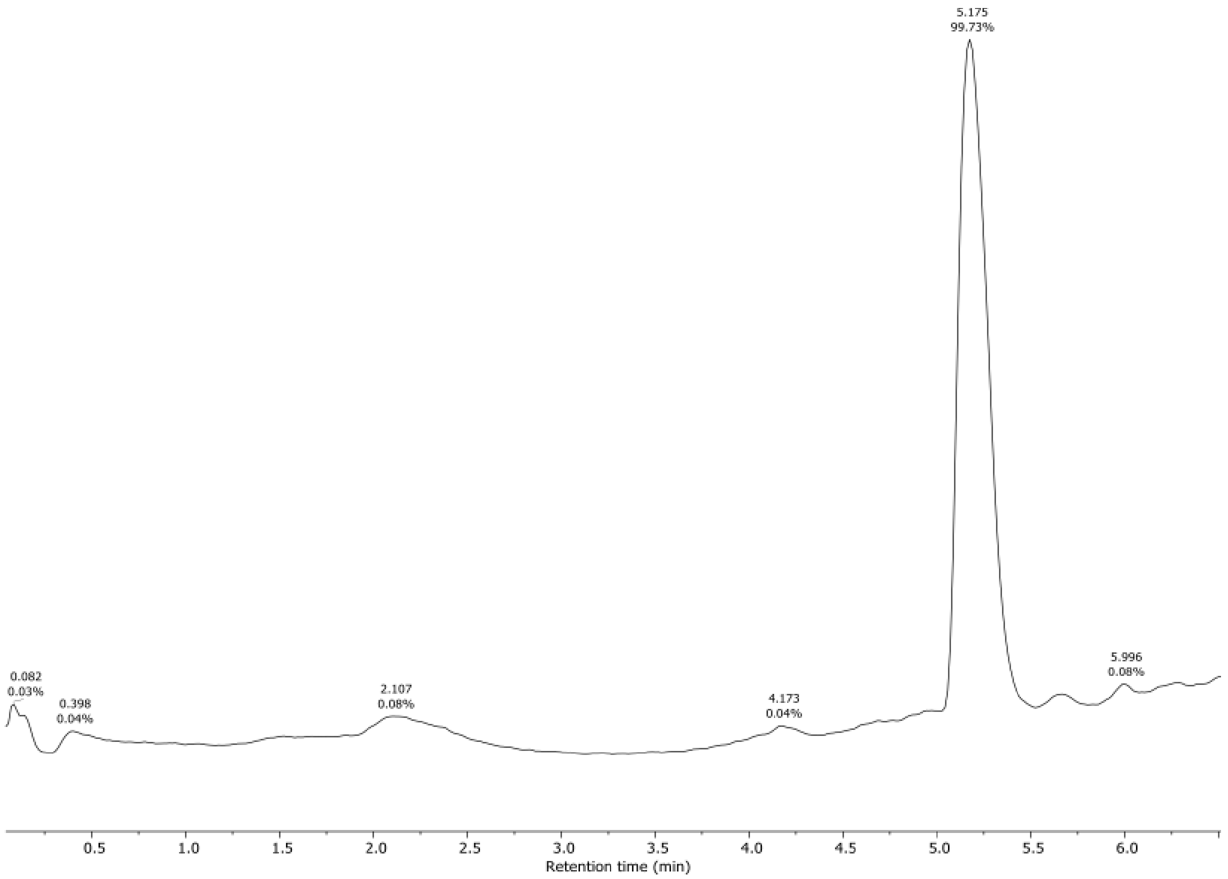

5.175
99.73%
0.082
0.03%
0.398
0.04%
2.107
0.08%
4.173
0.04%
5.996
0.08%
0.5 1.0 1.5 2.0 2.5 3.0 3.5 4.0 4.5 5.0 5.5 6.0
Retention time (min)


**HRMS**

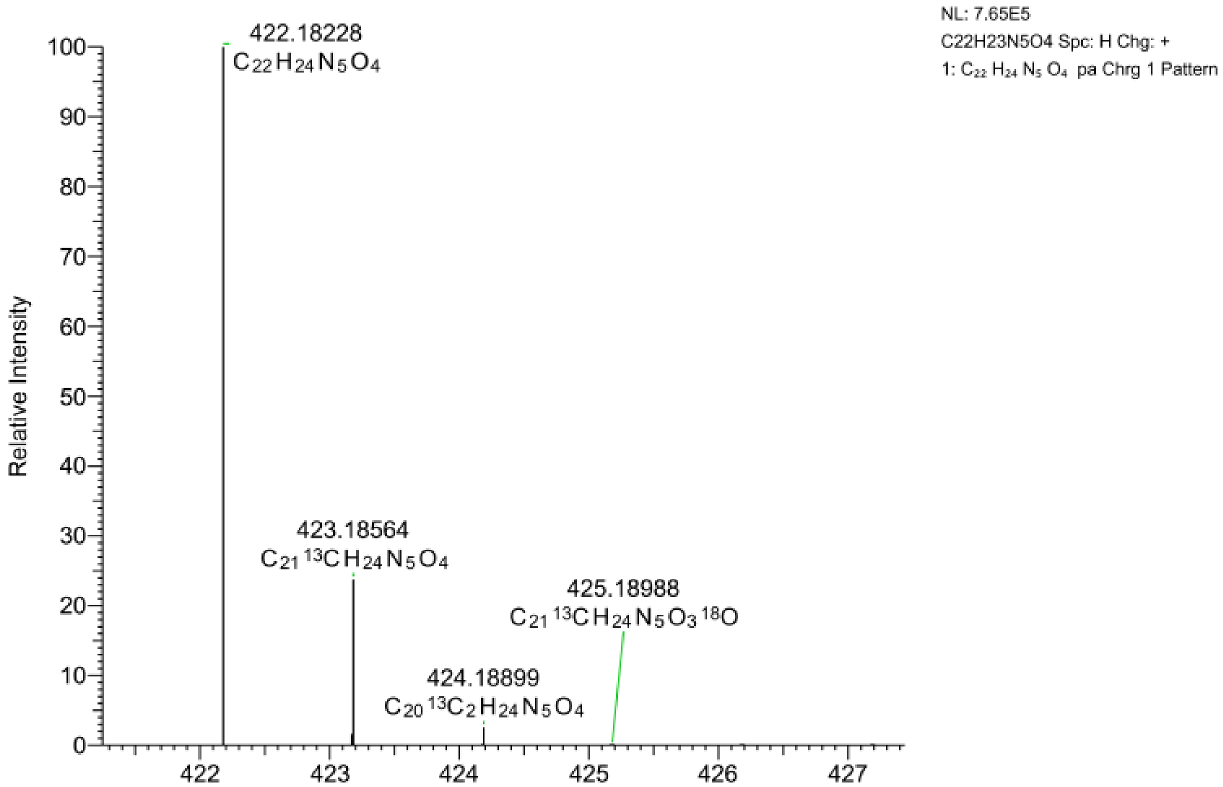

NL: 7.65E5
C22H23N5O4 Spc: H Chg: +
1: $C_{22}H_{24}N_5O_4$ pa Chrg 1 Pattern
422.18228
$C_{22}H_{24}N_5O_4$
423.18564
$C_{21}{}^{13}CH_{24}N_5O_4$
424.18899
$C_{20}{}^{13}C_2H_{24}N_5O_4$
425.18988
$C_{21}{}^{13}CH_{24}N_5O_3{}^{18}O$
Relative Intensity
m/z

***N*-(6-azidohexyl)-2-oxo-2-(2-phenyl-1H-indol-3-yl)acetamide**

**1H NMR**

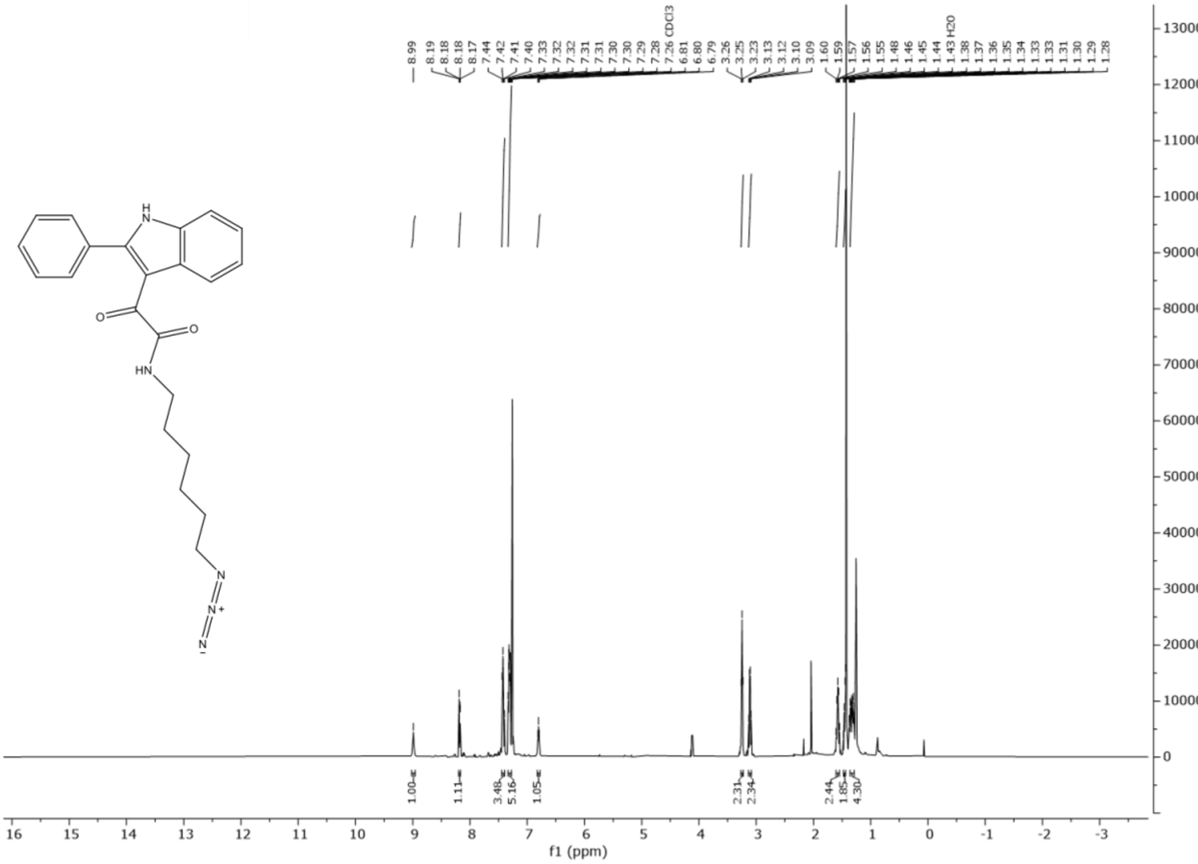


$^1$H-NMR (500MHz, MeOD, *J* Hz) δ ppm 10.21 (s, 1H), 8.17 (d, 1H, *J* = 8.00), 7.29 – 7.1 (m, 6H), 6.92 (dd, 3H, *J* = 7.45, 21.81), 3.66 – 3.56 (m, 6H), 3.43 (t, 2H, *J* = 5.10), 3.35 (t, 2H, *J* = 4.83), 3.22 (q, 2H, *J* = 5.12).

**13C NMR**

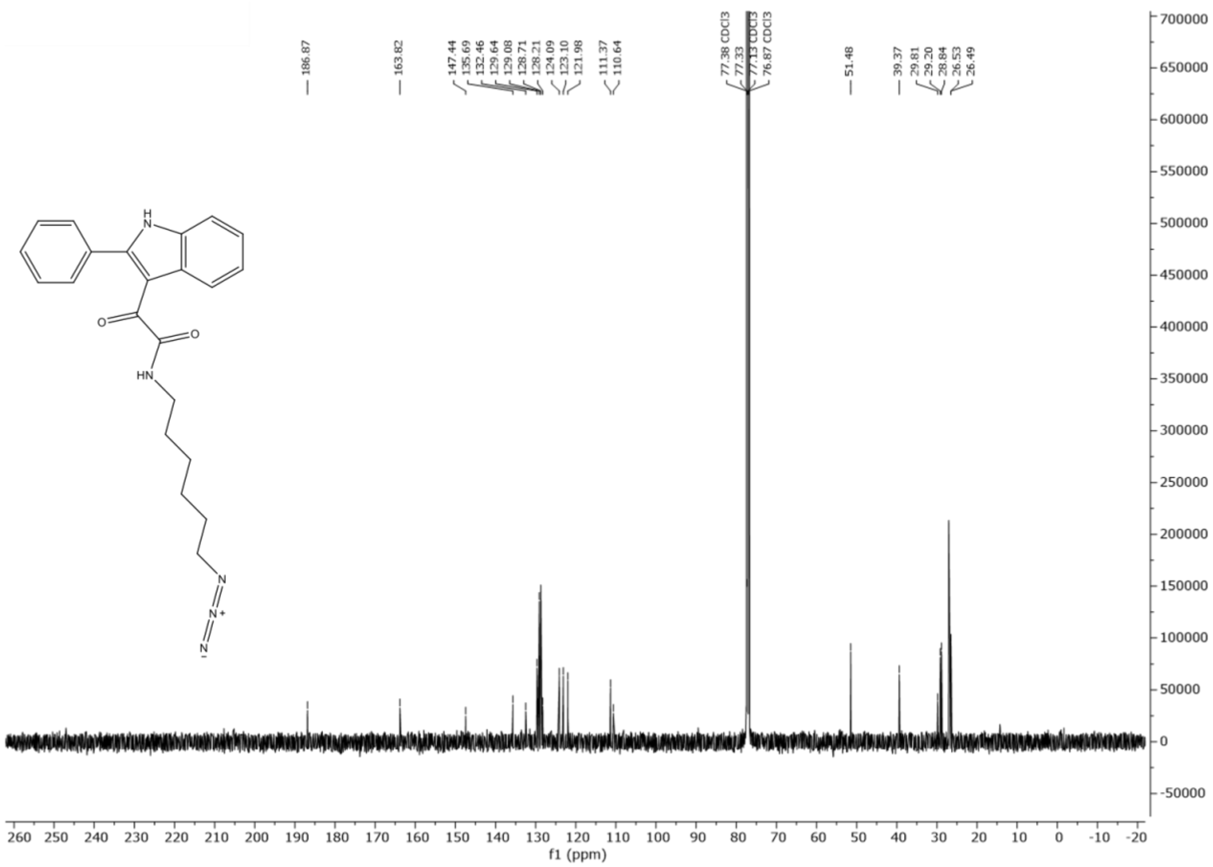


$^{13}$C-NMR (500MHz, MeOD) δ ppm 186.72, 171.31, 164.76, 147.76, 136.05, 132.05, 129.37, 129.11, 128.95, 128.51, 128.27, 123.78, 122.93, 121.57, 112.14, 110.17, 77.43, 77.17, 76.92, 70.71, 70.50, 70.22, 69.31, 60.53, 50.78, 39.24, 27.05, 21.14, 14.31.

**UV Chromatogram (LC-MS)**

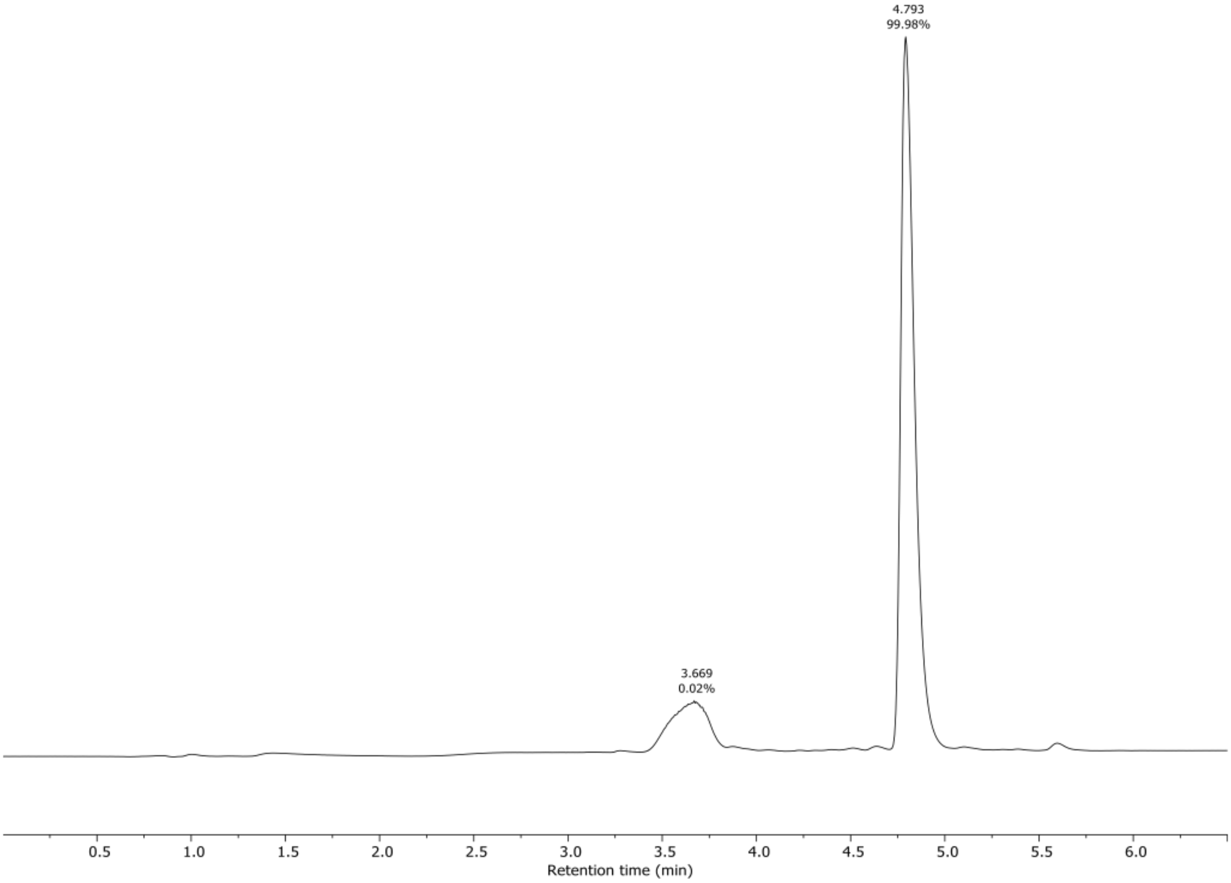


**HRMS**

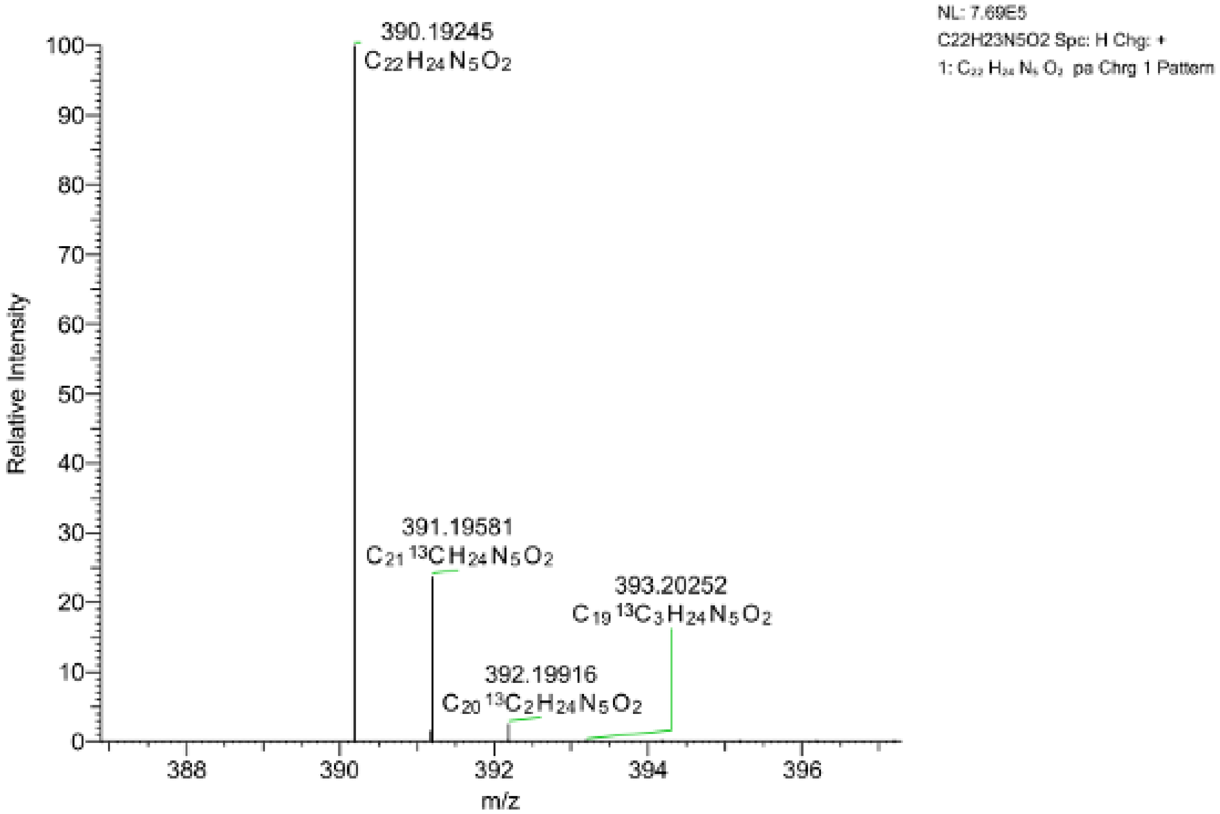


***(R)*-4-(1-amino-2-((2,4-difluorophenyl)amino)-2-oxoethyl)phenyl isobutyl carbonate**

**1H NMR**

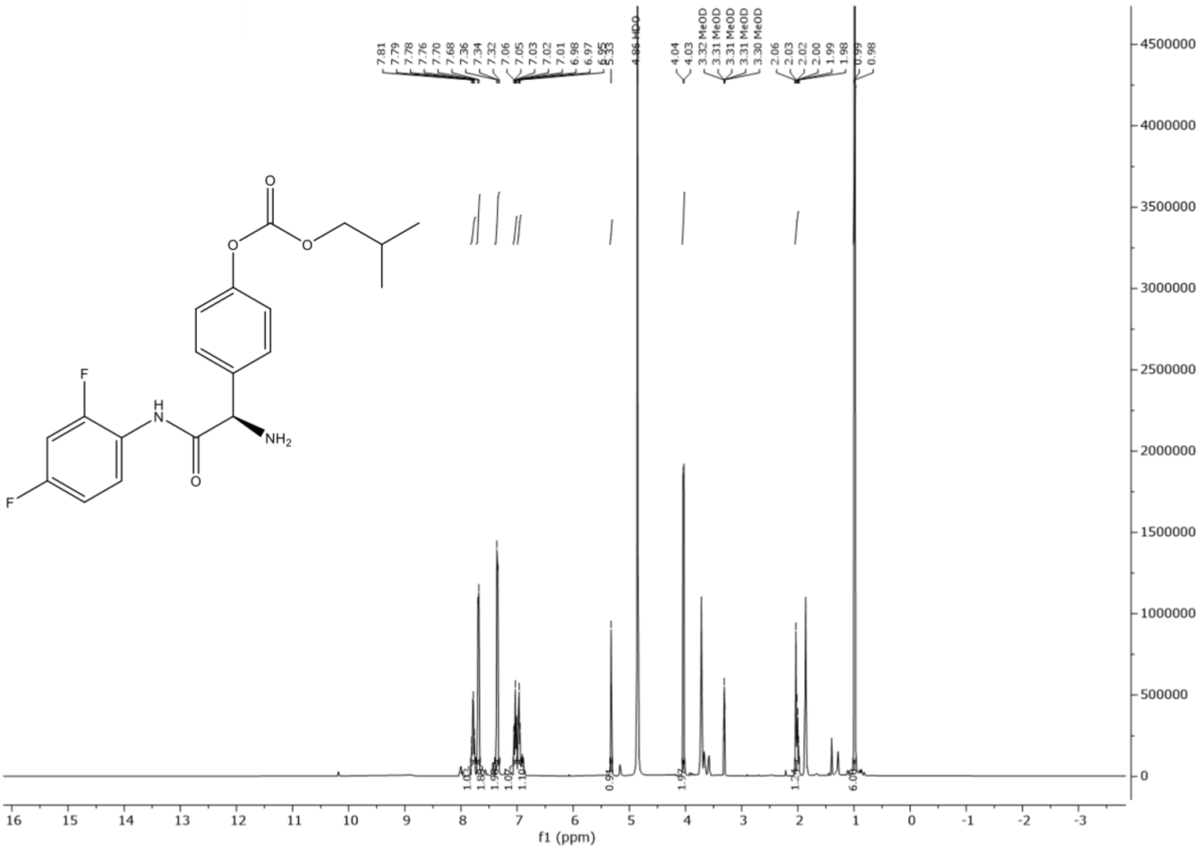

$^1$H-NMR (500MHz, $CDCl_3$, *J* Hz) δ ppm 7.828 – 7.736 (m, 1H), 7.689 (d, 2H, *J* = 8.223), 7.353 (d, 2H, *J* = 8.637), 7.070 – 7.006 (m, 1H), 6.975 (d, 1H, *J* = 9.240), 5.325 (s, 1H), 4.035 (d, 2H, *J* = 6.547), 2.053 – 1.990 (m, 1H), 0.988 (d, 6H, *J* = 6.744).

**13C NMR**

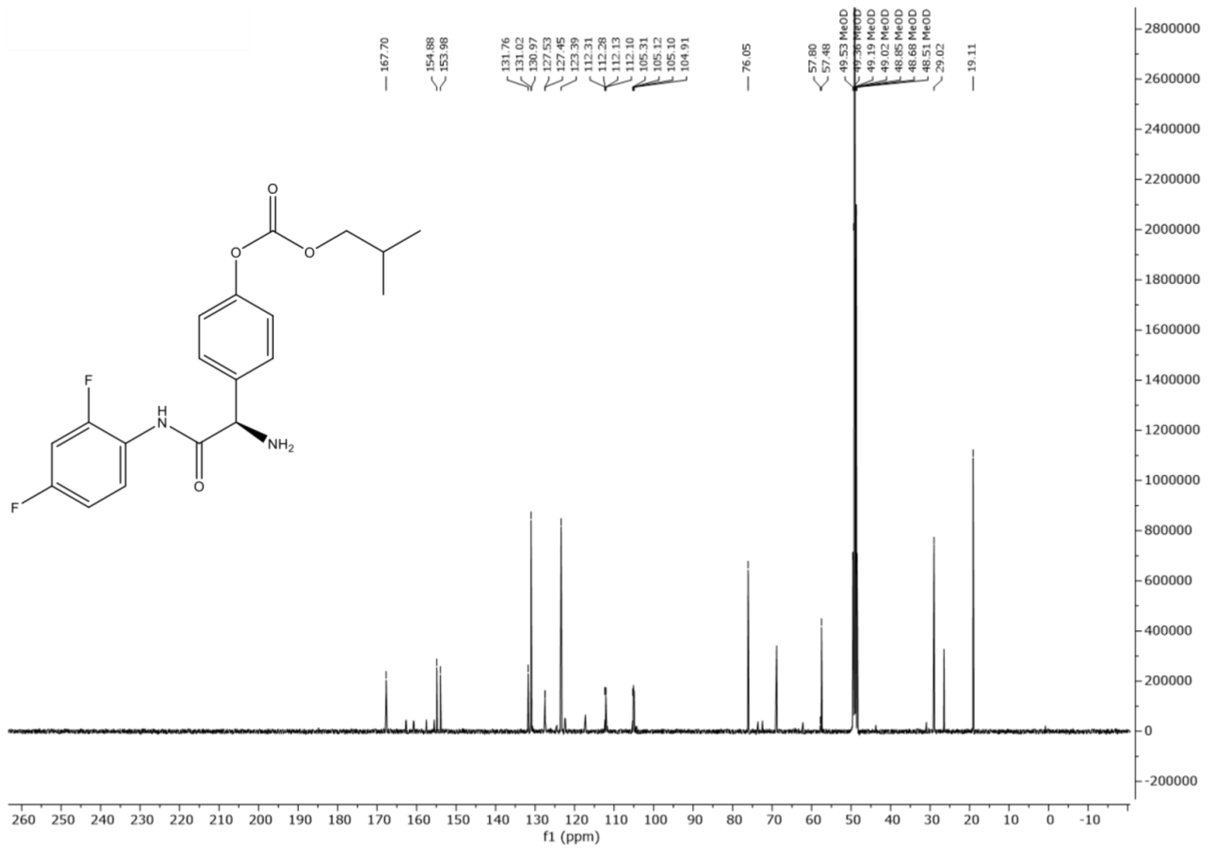


$^{13}$C-NMR (500MHz, $CDCl_3$) $\delta$ ppm 167.70, 154.88, 153.98, 131.76, 131.02, 130.97, 127.53, 127.45, 123.39, 112.31, 112.28, 112.13, 112.10, 105.31, 105.12, 105.10, 104.91, 76.05, 57.80, 57.48, 29.02, 19.11.

**UV Chromatogram (LC-MS)**

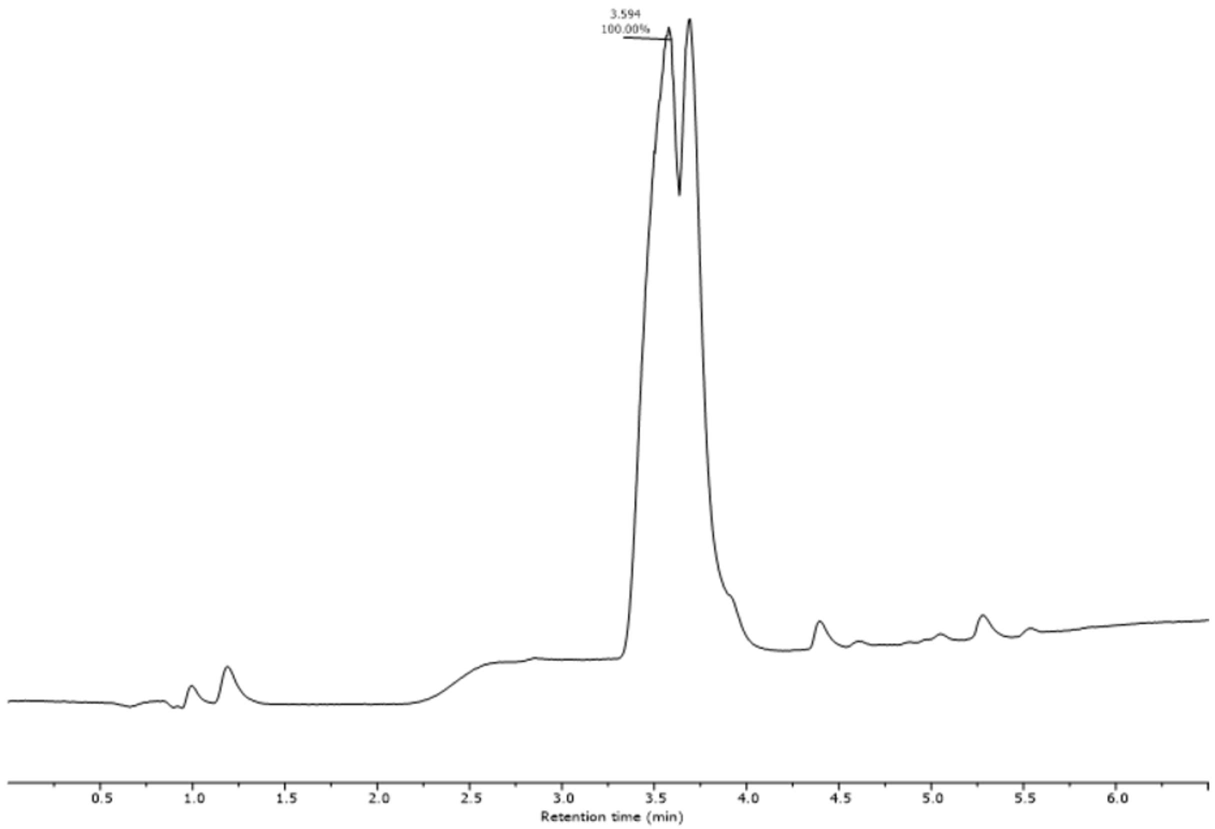


**HRMS**

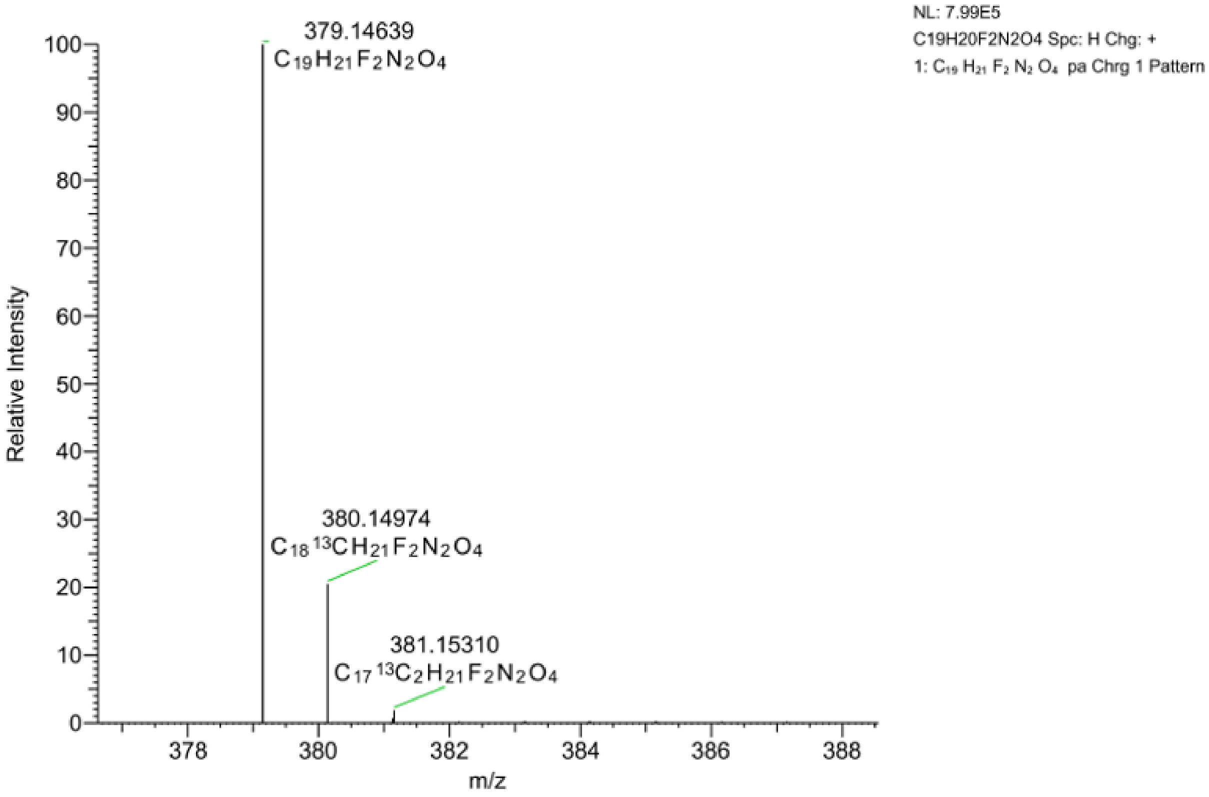


**(*R*)-2-(3-(3,5-bis(trifluoromethyl)phenyl)ureido)-N-(2,4-difluorophenyl)-2-(4-hydroxyphenyl)acetamide**

**1H NMR**

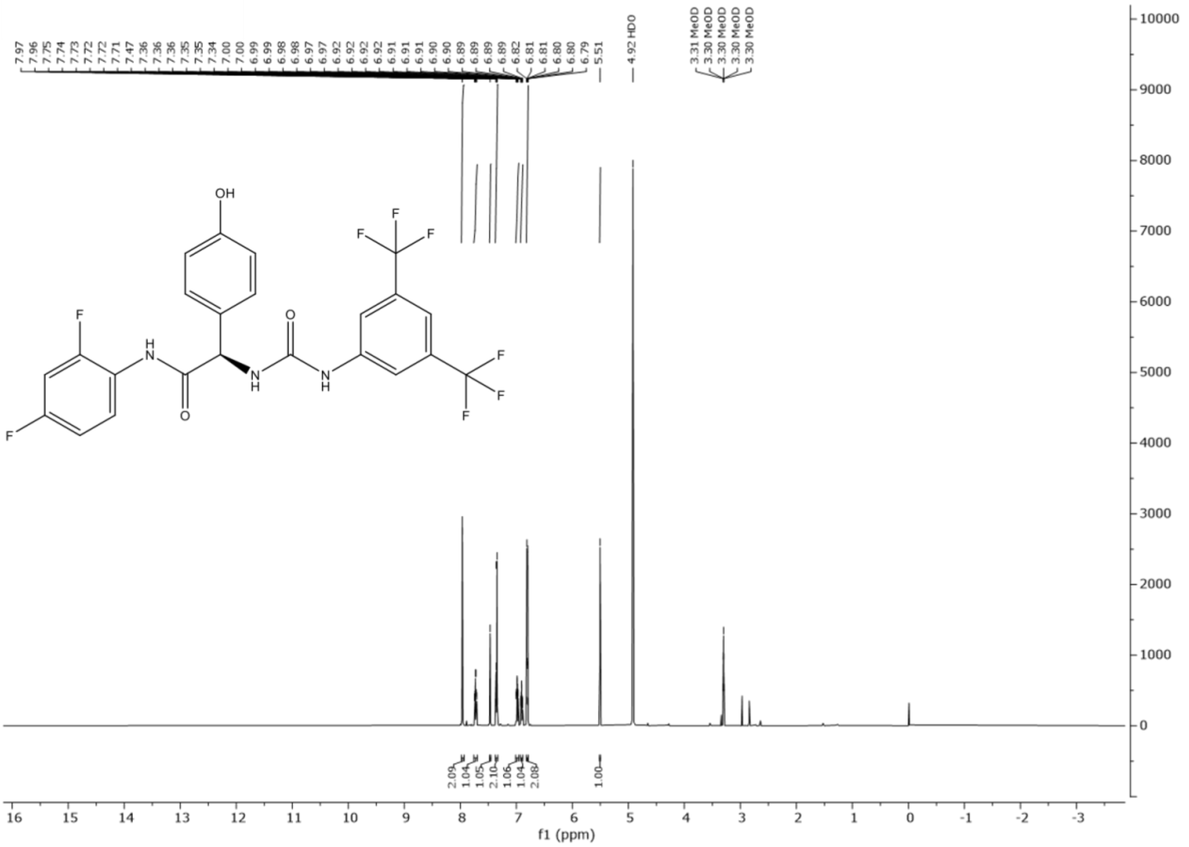


$^{1}$H-NMR (600MHz, MeOD, *J* Hz) $\delta$ ppm 7.99 (s, 2H), 7.73 (td, 1H, *J* = 8.89, 5.97), 7.47 (s, 1H), 7.36, (d, 1H, *J* = 8.55), 6.985 (ddd, 1H, *J* = 10.763, 8.763, 2.828), 6.905 (dddd, 1H, *J* = 9.245, 8.061, 2.855, 1.439), 6.82 (d, 1H, J = 8.55), 5.51 (s, 1H).

**13C NMR**

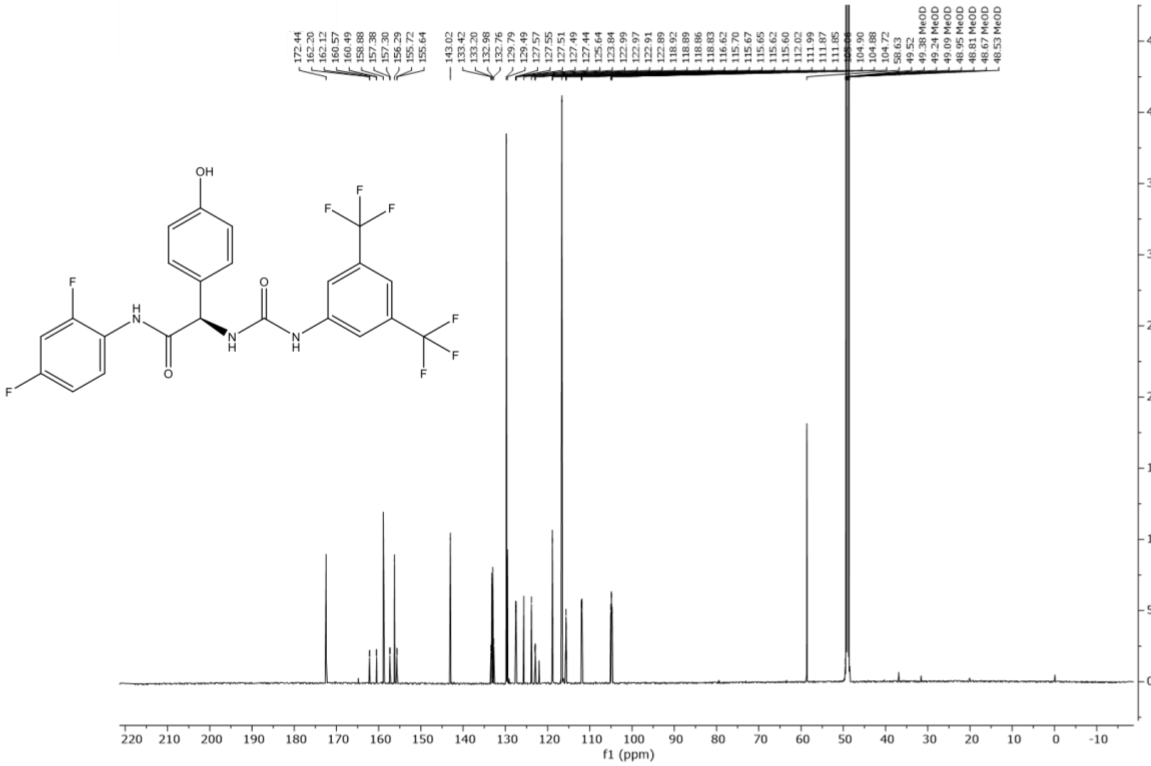


$^{13}C$-NMR (600MHz, MeOD) δ ppm 172.44, 162.20, 160.57, 158.88, 157.38, 156.29, 155.72, 143.02, 133.20, 129.79, 129.49, 127.51, 125.64, 123.84, 122.99, 118.86, 116.62, 115.62, 111.99, 104.90, 58.63.

**UV Chromatogram (LC-MS)**

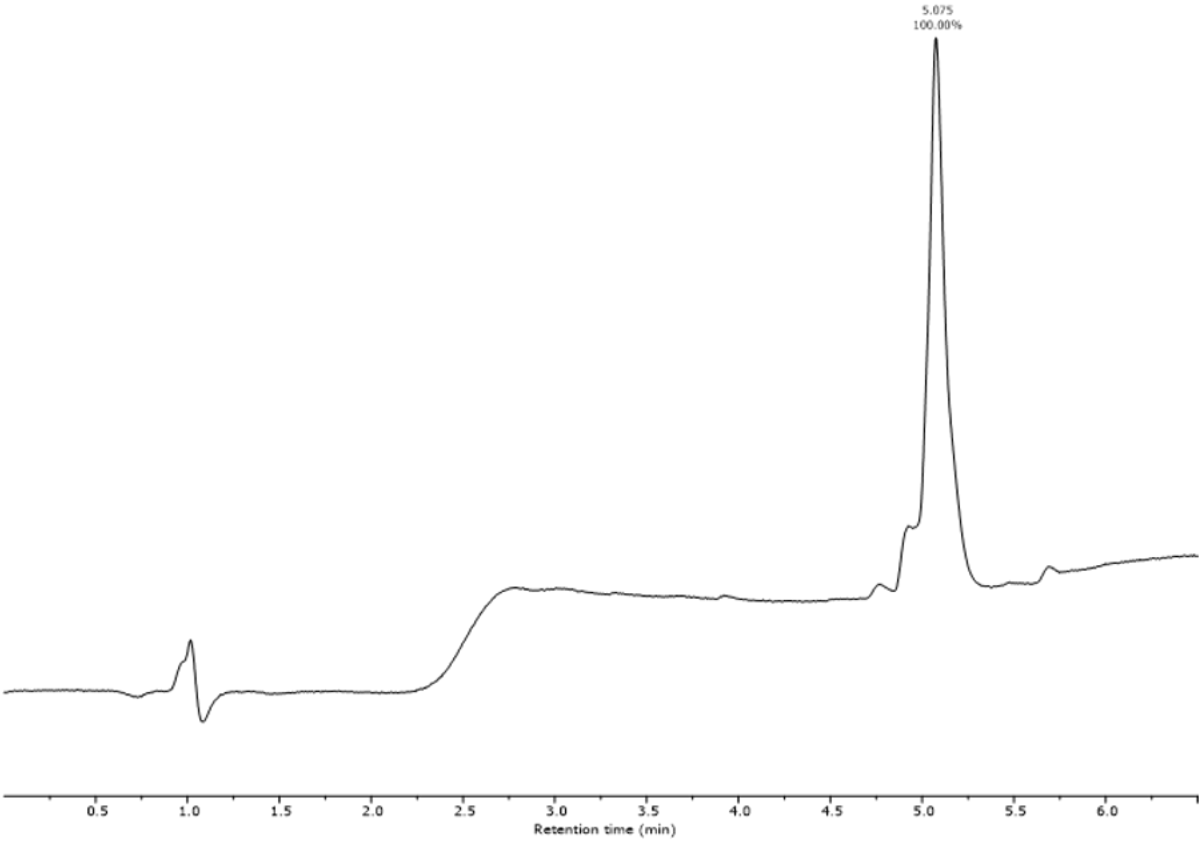
5.075
100.00%
0.5
1.0
1.5
2.0
2.5
3.0
3.5
4.0
4.5
5.0
5.5
6.0
Retention time (min)

**HRMS**

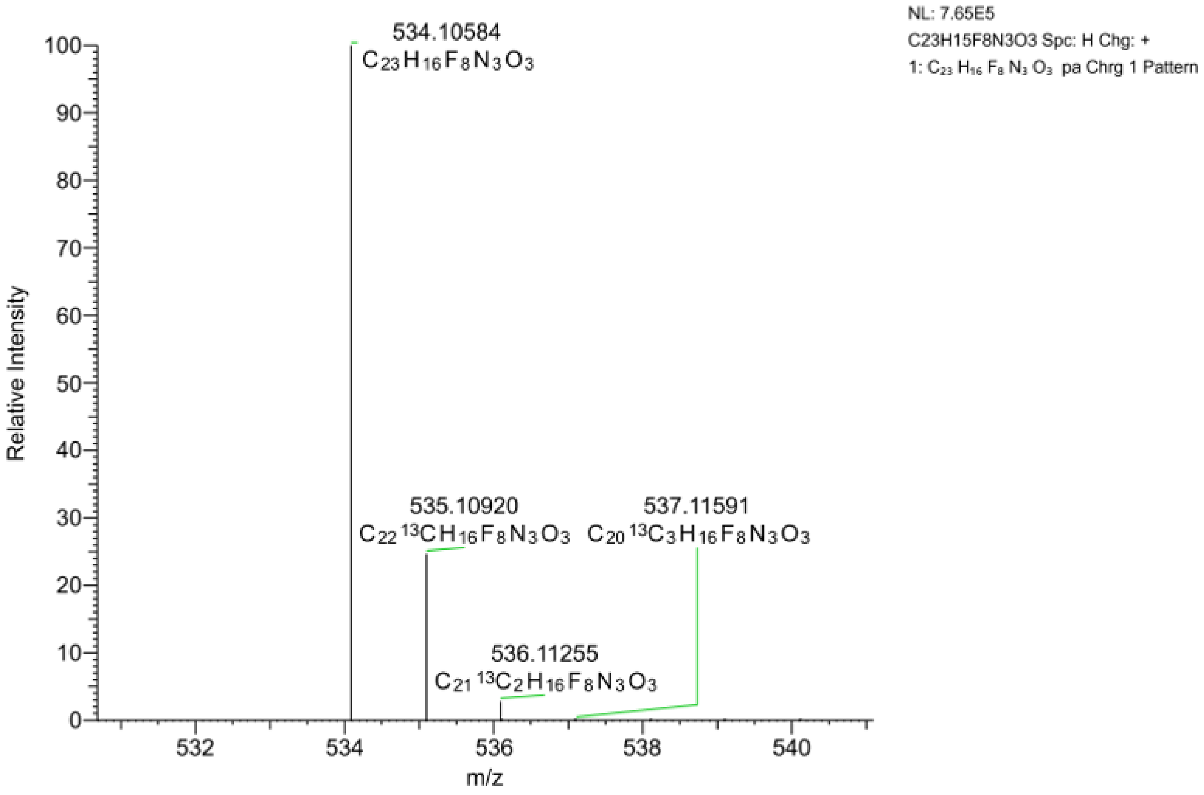
NL: 7.65E5
C23H15F8N3O3 Spc: H Chg: +
1: C23 H16 F8 N3 O3 pa Chrg 1 Pattern
534.10584
C23H16F8N3O3
535.10920
C22 13CH16F8N3O3
537.11591
C20 13C3H16F8N3O3
536.11255
C21 13C2H16F8N3O3
Relative Intensity
m/z
100
90
80
70
60
50
40
30
20
10
0
532
534
536
538
540

**(*R*)-2-(3-(3,5-bis(trifluoromethyl)phenyl)ureido)-*N*-(2,4-difluorophenyl)-2-(4-((1-(2-(2-(2-(2-oxo-2-(2-phenyl-1*H*-indol-3-yl)acetamido)ethoxy)ethoxy)ethyl)-1*H*-1,2,3-triazol-4-yl)methoxy)phenyl)acetamide [NZ-65]**

**1H NMR**

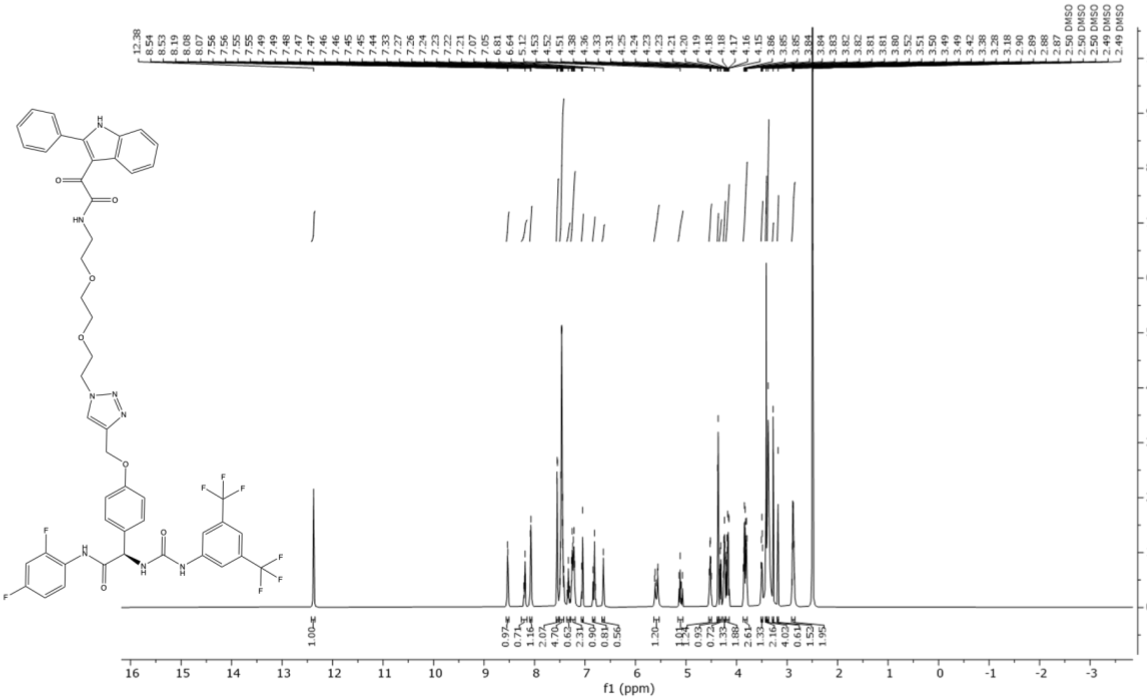


$^{1}$H NMR (600 MHz, DMSO-$d_6$, *J* Hz) δ 12.38 (s, 1H), 8.53 (d, 1H, J = 5.4), 8.19 (q, 1H, J = 7.1), 8.08 (d, 1H, J = 7.4), 7.55 (dd, 2H, J = 4.2, 2.1), 7.52 – 7.41 (m, 5H), 7.32 (q, 1H, J = 9.5), 7.24 (dt, 2H, J = 21.8, 7.4), 7.06 (d, 1H, J = 10.6), 6.80 (d, 1H, J = 12.6), 6.64 (s, 1H), 5.63 – 5.54 (m, 1H), 5.11 (dd, 1H, J = 27.9, 12.7), 4.52 (p, 1H, J = 7.3), 4.37 (d, 1H, J = 8.6), 4.34 – 4.30 (m, 1H), 4.25 (q, 1H, J = 6.1), 4.18 (ddd, 2H, J = 19.6, 9.2, 6.4), 3.83 (tdd, 3H, J = 12.9, 9.5, 5.2), 3.50 (dt, 1H, J = 7.4, 3.6), 3.42 (s, 2H), 3.38 (s, 4H), 3.28 (s, 1H), 3.18 (s, 2H), 2.88 (h, 2H, J = 6.0).

**13C NMR**

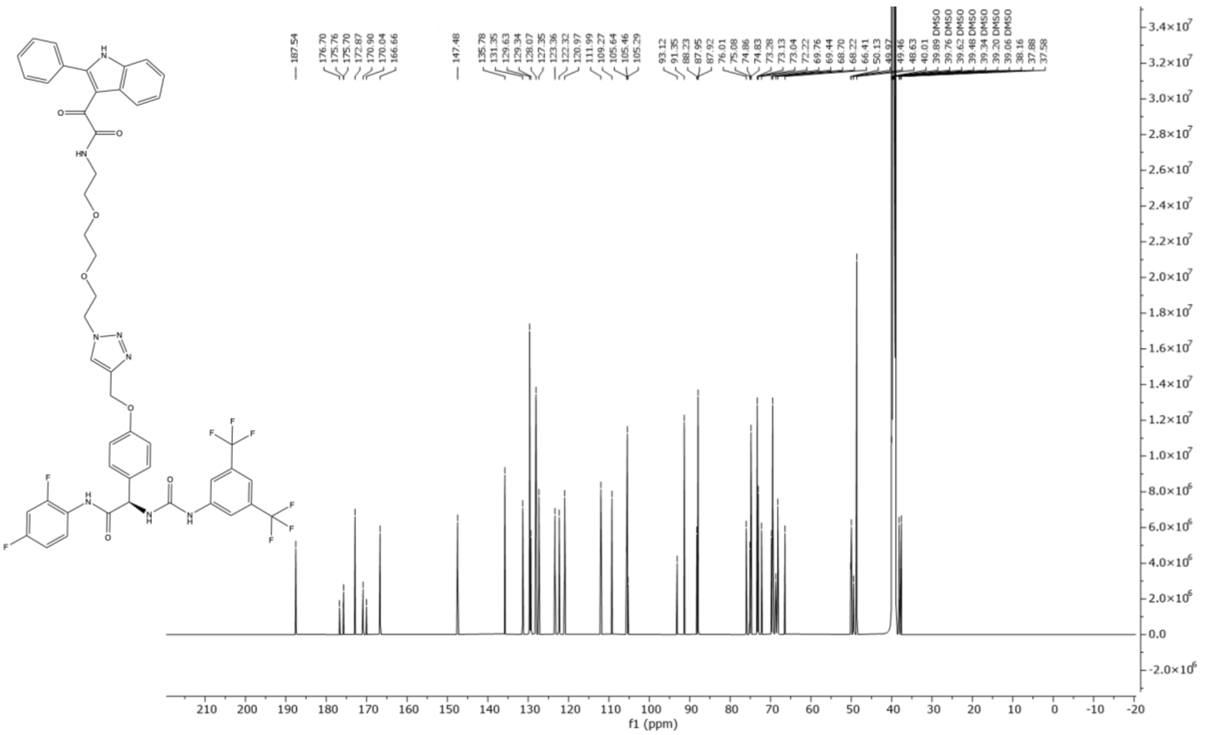


$^{13}C$ NMR (600 MHz, DMSO-$d_6$) δ ppm 187.54, 176.70, 175.76, 175.70, 172.87, 170.90, 170.04, 166.66, 147.48, 135.78, 131.35, 129.63, 129.34, 128.07, 127.35, 123.36, 122.32, 120.97, 111.99, 109.27, 105.64, 105.46, 105.29, 93.12, 91.35, 88.23, 87.95, 87.92, 76.01, 75.08, 74.86, 74.83, 73.28, 73.13, 73.04, 72.22, 69.76, 69.44, 68.70, 68.22, 66.41, 50.13, 49.97, 49.46, 48.63, 40.01, 38.16, 37.88, 37.58.

**UV Chromatogram (LC-MS)**

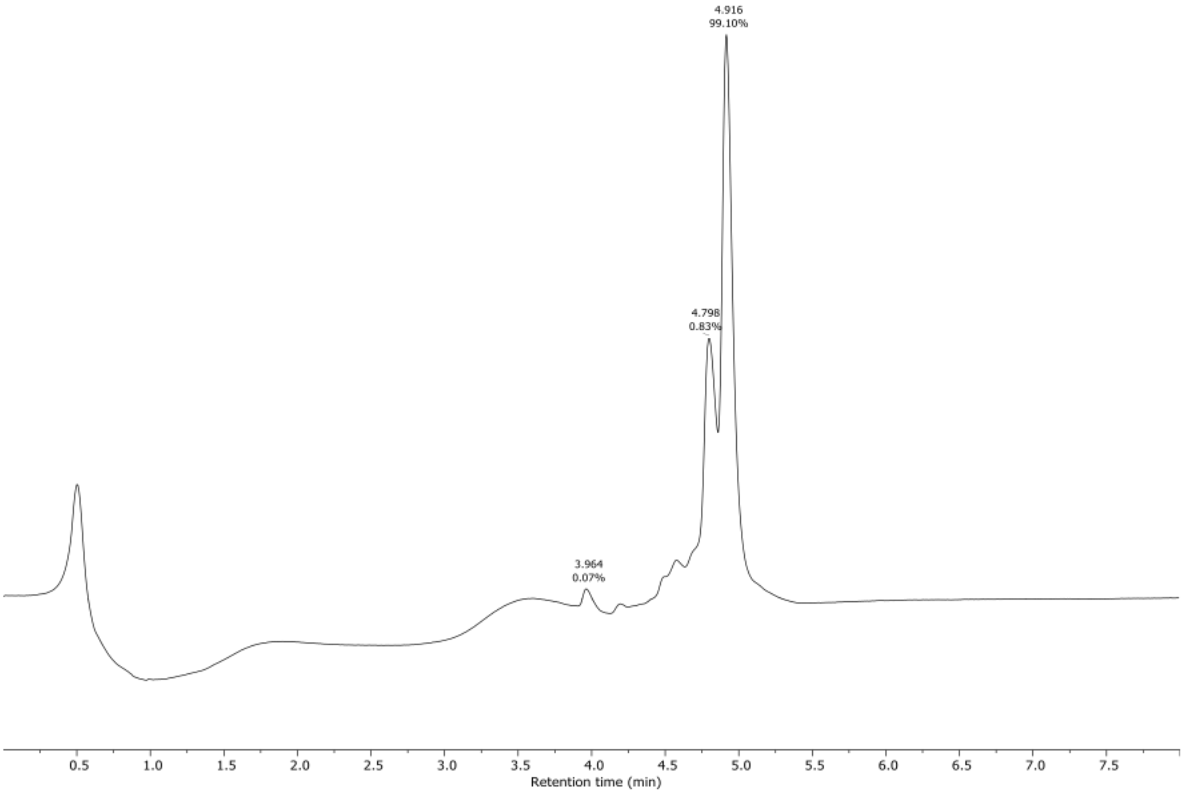


**HRMS**

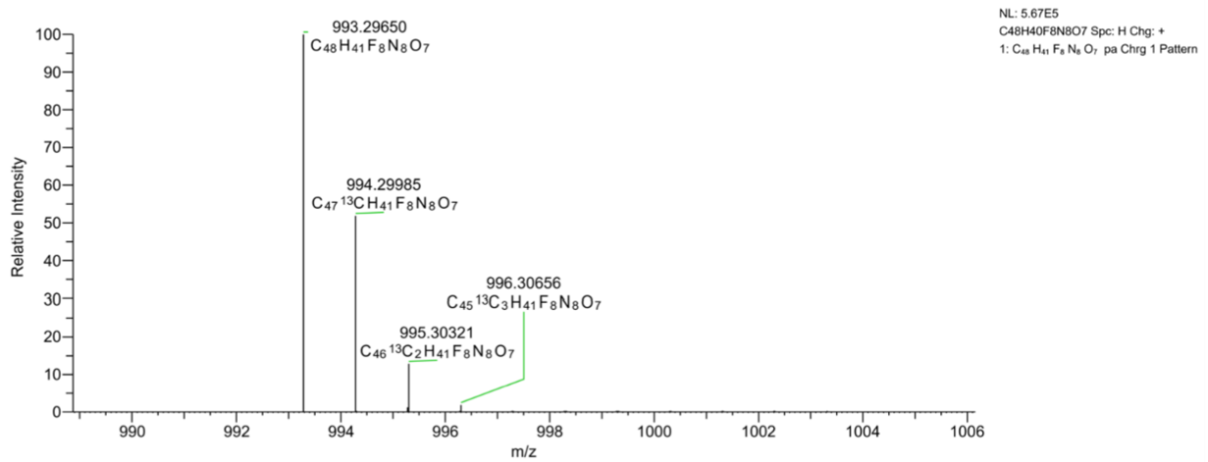


**(*R*)-2-(3-(3,5-bis(trifluoromethyl)phenyl)ureido)-*N*-(2,4-difluorophenyl)-2-(4-((1-(6-(2-oxo-2-(2-phenyl-1*H*-indol-3-yl)acetamido)hexyl)-1*H*-1,2,3-triazol-4-yl)methoxy)phenyl)acetamide [NZ-66]**

**1H NMR**

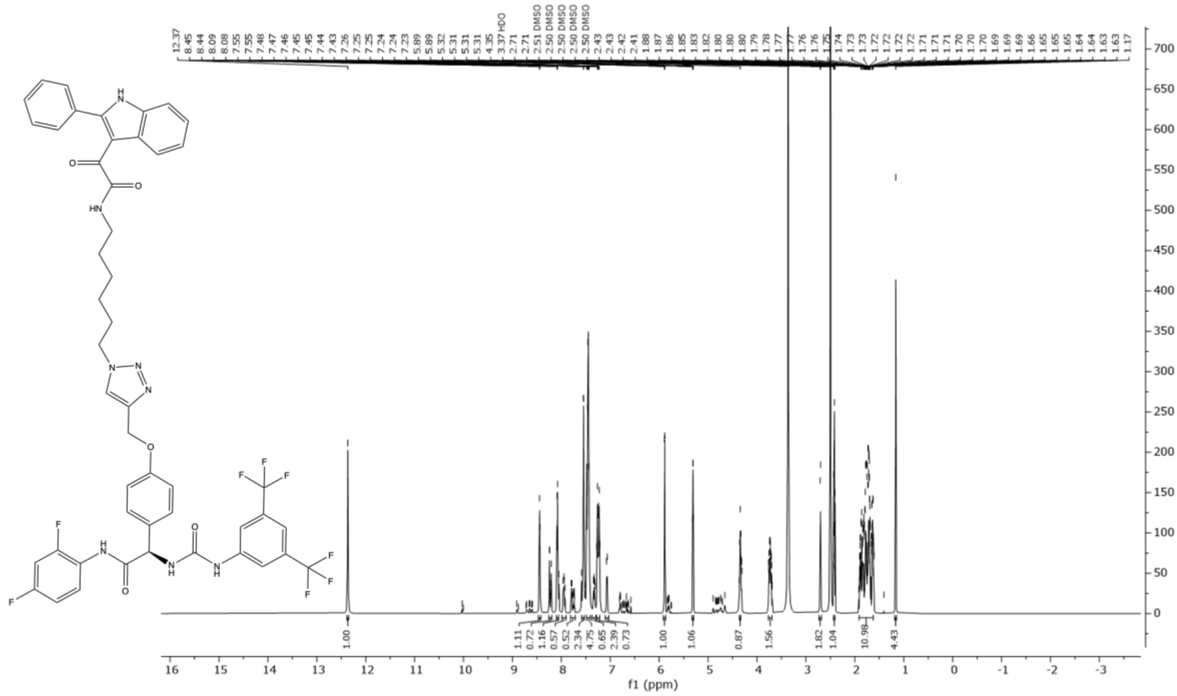


$^{1}$H NMR (600 MHz, DMSO-$d_6$, *J* Hz) δ 12.37 (s, 1H), 8.44 (d, 1H, J = 5.7), 8.23 (dd, 1H, J = 19.1, 4.1), 8.09 (d, 1H, J = 7.7), 7.95 (dd, 1H, J = 12.9, 10.5), 7.81 – 7.71 (m, 1H), 7.55 (d, 2H, J = 4.1), 7.49 – 7.41 (m, 5H), 7.33 (ddd, 1H, J = 13.9, 8.0, 3.1), 7.25 (dt, 2H, J = 21.4, 7.5), 7.07 (d, 1H, J = 8.8), 5.89 (d, 1H, J = 4.7), 5.31 (q, 1H, J = 3.0), 4.35 (t, 1H, J = 7.1), 3.73 (dddt, 2H, J = 15.4, 7.8, 5.6, 3.6), 2.71 (s, 2H), 2.42 (t, 1H, J = 8.1), 1.91 – 1.62 (m, 11H), 1.17 (s, 4H).

**13C NMR**

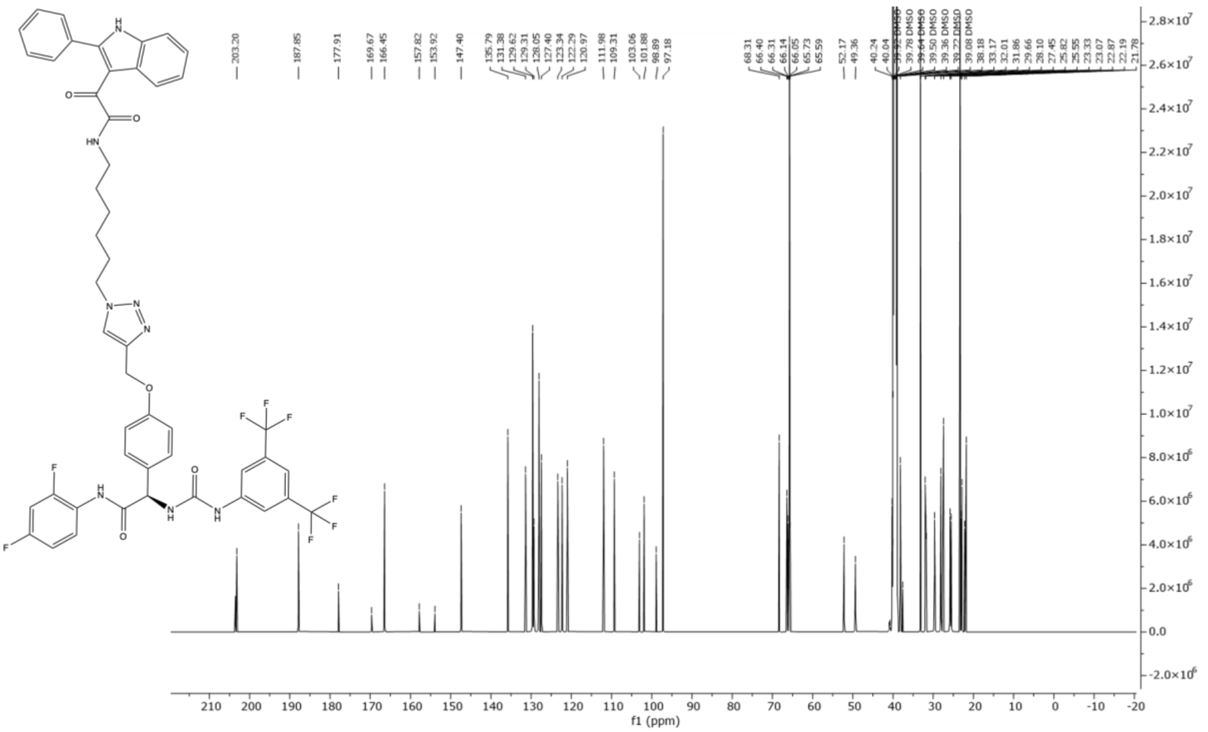


$^{13}C$ NMR (600 MHz, DMSO-$d_6$) δ ppm 203.53, 203.20, 187.85, 177.91, 169.67, 166.45, 157.82, 153.92, 147.40, 135.79, 131.38, 129.62, 129.31, 128.05, 127.40, 123.34, 122.29, 120.97, 111.98, 109.31, 103.06, 101.88, 98.89, 97.18, 68.31, 66.40, 66.31, 66.14, 66.05, 65.73, 65.58, 52.17, 49.36, 40.96, 40.74, 40.24, 40.10, 40.04, 38.18, 37.58, 33.17, 32.01, 31.86, 31.75, 29.66, 28.10, 27.45, 25.82, 25.55, 23.33, 23.07, 23.00, 22.87, 22.19, 21.78.

**UV Chromatogram (LC-MS)**

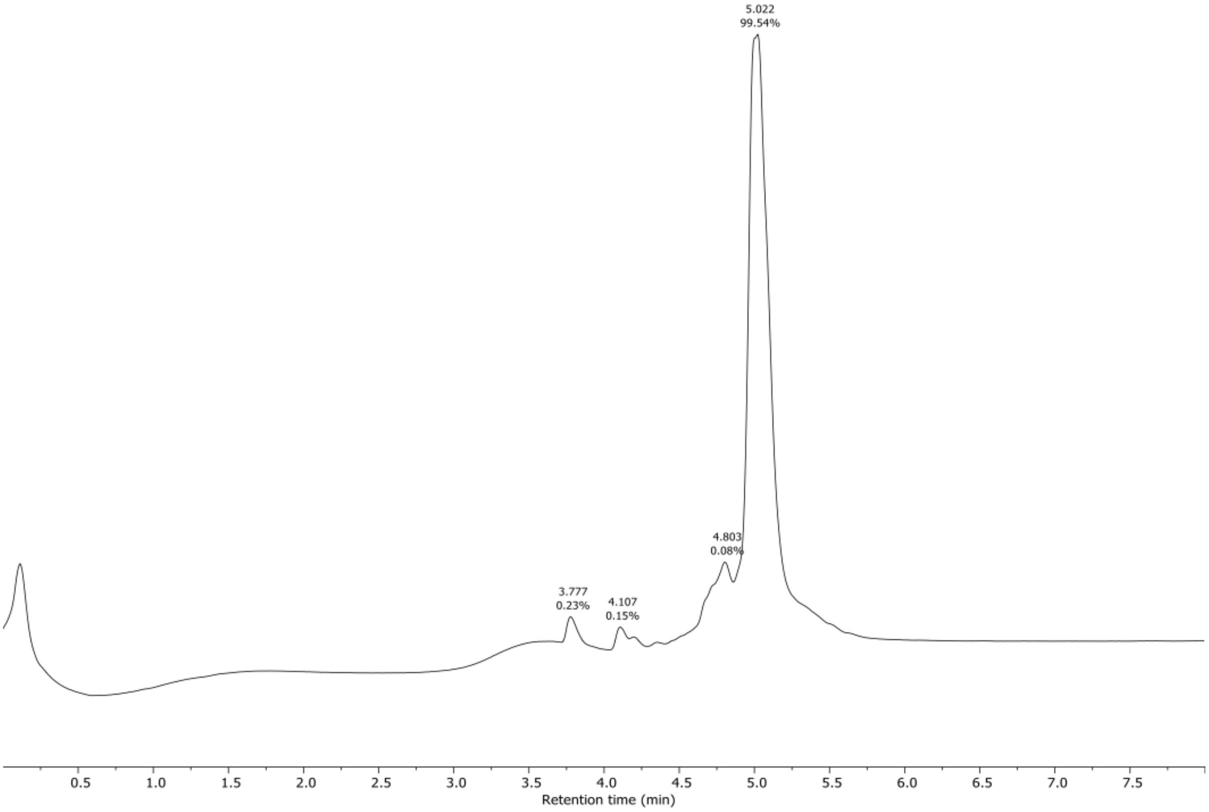

**HRMS**

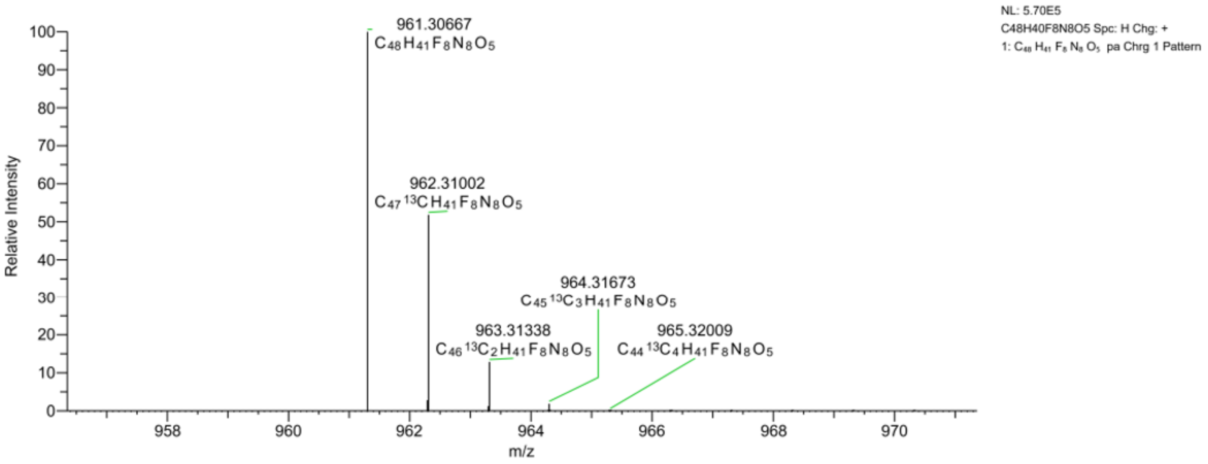

NL: 5.70E5
C48H40F8N8O5 Spc: H Chg: +
1: C48 H41 F8 N8 O5 pa Chrg 1 Pattern
961.30667
C48H41F8N8O5
962.31002
C47 13C H41F8N8O5
963.31338
C46 13C2 H41F8N8O5
964.31673
C45 13C3 H41F8N8O5
965.32009
C44 13C4 H41F8N8O5
Relative Intensity
m/z
